# Barcode Sequencing and a High-throughput Assay for Chronological Lifespan Uncover Ageing-associated Genes in Fission Yeast

**DOI:** 10.1101/2021.03.04.433786

**Authors:** Catalina A. Romila, StJohn Townsend, Michal Malecki, Stephan Kamrad, María Rodríguez-López, Olivia Hillson, Cristina Cotobal, Markus Ralser, Jürg Bähler

## Abstract

Ageing-related processes are largely conserved, with simple organisms remaining the main platform to discover and dissect new ageing-associated genes. Yeasts provide potent model systems to study cellular ageing owing their amenability to systematic functional assays under controlled conditions. Even with yeast cells, however, ageing assays can be laborious and resource-intensive. Here we present improved experimental and computational methods to study chronological lifespan in *Schizosaccharomyces pombe*. We decoded the barcodes for 3206 mutants of the latest gene-deletion library, enabling the parallel profiling of ∼700 additional mutants compared to previous screens. We then applied a refined method of barcode sequencing (Bar-seq), addressing technical and statistical issues raised by persisting DNA in dead cells and sampling bottlenecks in aged cultures, to screen for mutants showing altered lifespan during stationary phase. This screen identified 341 long-lived mutants and 1246 short-lived mutants which point to many previously unknown ageing-associated genes, including 51 conserved but entirely uncharacterized genes. The ageing-associated genes showed coherent enrichments in processes also associated with human ageing, particularly with respect to ageing in non-proliferative brain cells. We also developed an automated colony-forming unit assay for chronological lifespan to facilitate medium- to high-throughput ageing studies by saving time and resources compared to the traditional assay. Results from the Bar-seq screen showed good agreement with this new assay, validating 33 genes not previously associated with cellular ageing. This study provides an effective methodological platform and identifies many new ageing-associated genes as a framework for analysing cellular ageing in yeast and beyond.

## INTRODUCTION

Ageing is a multifactorial process leading to a gradual decline in biological function over time [1–3]. Old age is the main risk factor for several complex diseases including diabetes, neurodegeneration, cardiovascular disease and cancer. The study of specific disease mechanisms has long been a focus of biomedical research, but it is also imperative to consider fundamental aspects of ageing as a vital part of the problem and to explore ways to slow its effects. Ageing research has been galvanised by the discovery of lifespan-extending mutations in worms [4], with subsequent research identifying hundreds of ageing-related genes in various model systems [1, 5–7]. Age-related decline is plastic, with multiple genetic factors and biological processes contributing to lifespan and ageing. Owing to its complexity, genetic and genomic research on ageing in simple model organisms remains vital to discover all proteins and processes affecting lifespan [8]. Ageing experiments are often laborious and resource-consuming, especially in vertebrate models which can live for several years. Moreover, ageing experiments typically require large sample sizes owing to poor experimental reproducibility and substantial phenotypic variability in lifespan even amongst genetically identical individuals [9, 10]. This situation highlights the need for tractable experimental approaches which facilitate systematic and well-controlled lifespan assays.

Yeast cells are well-established as a system to carry out systematic, genome-scale studies: they are relatively simple and can be cultured under tightly controlled conditions in parallelised experimental platforms [11]. The budding yeast, *Saccharomyces cerevisiae*, and the distantly related fission yeast, *Schizosaccharomyces pombe*, are also established ageing models. The processes affecting longevity are remarkably well conserved from yeast to human, including both genetic factors, such as the TORC1 nutrient-sensing pathway, and environmental factors, such as dietary restriction [2, 12, 13]. Fission yeast has a well-annotated genome encoding 5064 proteins, about 70% of which have identifiable human orthologs [14]. It did not undergo any genome duplication and features lower gene redundancy, with mutants thus being more likely to show phenotypes. In addition, ∼80% of all *S. pombe* genes are expressed under standard growth conditions [15], which greatly facilitates functional studies.

We and others have explored the effects of nutrient limitation, signalling pathways, and gene deletions on the chronological lifespan (CLS) of *S. pombe* cells, and several ageing-associated proteins have been identified [16–25]. CLS is defined as the time a cell remains viable in a non-dividing state, which mirrors ageing of post-mitotic or quiescent cells in multi-cellular organisms [2, 13]. CLS is typically measured in stationary phase cultures following glucose exhaustion, where *S. pombe* cells mostly arrest in the G2 phase of the cell cycle and die within a few days. Chronological ageing can be induced by depleting cells of other nutrients such as nitrogen, where *S. pombe* cells reversibly arrest in a G0-like state and survive for many weeks [19], or even by physically restricting cells such that they cannot divide [26].

CLS is traditionally measured by counting the number of colony-forming units (CFUs) grown from ageing cell cultures after spreading cell aliquots on solid agar plates. The cell aliquots need to be serially diluted and plated at different concentrations to quantify the number of CFUs. Hence, measuring CLS via CFUs is error-prone, laborious and resource-intense, and it does not scale to larger studies. Several alternatives to the traditional CFU assay have been proposed: cells are cultured in a high-throughput format and CLS is determined via an alternative approach, such as measuring the proportion of cells stained with a viability dye using a flow cytometer [27] or fluorescent plate reader [28, 29], inoculating re-growth cultures and measuring optical density as a proxy for the number of viable cells in the inoculum [30], or competitively ageing fluorescently-tagged strains and measuring relative fluorescence of re-growth cultures in a plate reader [31]. Alternatively, genome-wide collections of non-essential deletion mutants can be pooled and aged competitively, where mutants with altered CLS are detected by quantifying the abundance of specific DNA barcodes associated with each mutant. This can be done via DNA microarrays [32] or next-generation sequencing, known as barcode sequencing, or Bar-seq [33, 34]. We have applied Bar-seq to screen an early version of the *S. pombe* deletion library for lifespan mutants during long-term quiescence [19]. Whilst large-scale screens have identified many ageing-related genes, there is a remarkably poor overlap between screens [31]. This irreproducibility could partly reflect experimental and analytical differences, but may also have biological origins. The genetic factors which determine CLS can differ depending on environmental conditions [27], with subtle changes in culture conditions altering the genetic basis of lifespan [35]. The gene-environment interactions uncovered in yeast CLS screens indicate that the genetics of lifespan is context-dependent. Understanding the genetics of lifespan as a function of environmental, physiological or pharmacological perturbations will help to develop a comprehensive view of ageing in yeast and beyond. Hence, there is a need for tractable experimental and analytical approaches which facilitate high-throughput, systematic and robust identification of determinants of CLS.

In this work, we present two approaches to study CLS for medium- to high-throughput applications. We apply a refined method for Bar-seq, along with a tailored analysis pipeline, to identify mutants showing altered CLS under glucose exhaustion during stationary phase. We also present a novel medium-throughput CFU assay that can be largely automated by robotics, which we use to validate the lifespan of mutants from the Bar-seq screen. This work provides a toolbox for systematic ageing studies at various experimental scales and serves as a basis to better understand the genetic basis and cellular mechanisms of ageing.

## RESULTS & DISCUSSION

### Barcode decoding of latest *S. pombe* deletion-mutant library

We first needed to decode the two unique barcode sequences (UpTag and DnTag) associated with each mutant as this information was only available for earlier versions of the deletion library [19, 33]. We could decode barcodes for 3206 gene-deletion mutants (94% of all mutants in this library), including 3011 mutants decoded for both UpTag and DnTag as well as 96 and 99 mutants decoded for UpTag or DnTag, respectively (Table S1; Materials and Methods). Reassuringly, the sequence counts for the UpTag and DnTag barcodes strongly correlated with each other (Figure S1A). As expected, most of the decoded barcodes were 20 nucleotides long, with a range of 14-22 nucleotides (Figure S1B), as reported [36]. As part of the decoding process, we visually confirmed the barcodes using an in-house genome browser (Figure S1C). This effort captured proportionately more barcode sequences than for previous library versions, which include 2560 decoded mutants (90% of library; [33]) and 2473 decoded mutants (82% of library; [19]). The effective decoding, along with the increased size of the latest deletion library, allowed us to interrogate a substantially higher number of deletion mutants by Bar-seq than in previous screens.

### Bar-seq screen for chronological lifespan of stationary-phase mutants

We developed a CLS screen to identify long- and short-lived deletion mutants by letting a mutant pool compete for survival in stationary phase followed by Bar-seq to determine the relative barcode abundance for each mutant as a function of age (Figure 1A). We carried out analyses to test the experimental design of the screen. Since Bar-seq relies on barcode sequences, any persisting DNA from dead cells may produce misleading results. We tested for this potential bias by chronologically ageing stationary-phase mutant pools for 6 days, with daily measurements of CFUs and DNA levels. The results indicated that DNA can indeed remain intact for several days following cell death (or loss of proliferative potential) (Figure 1B). This finding confirmed the DNA bias presumed in previous competition-based screens [19, 32, 37, 38]. To account for this bias, besides directly sampling non-dividing cells of the mutant pools, we also put aliquots of these pools in fresh medium and re-grew them to stationary phase (Figure 1A). This approach is similar to re-growth applied for another competition-based CLS screens [32]. We carried out three biological repeats of the chronological ageing and re-growth experiment using independent deletion-mutant pools, along with two wild-type control strains. We collected samples at 7 timepoints over 11 days (Figure 1A). In two independent repeats of the screen, the CLS of the mutant pools were slightly shorter than those of the wild-type strains (Figure 1C).

**Figure 1:**
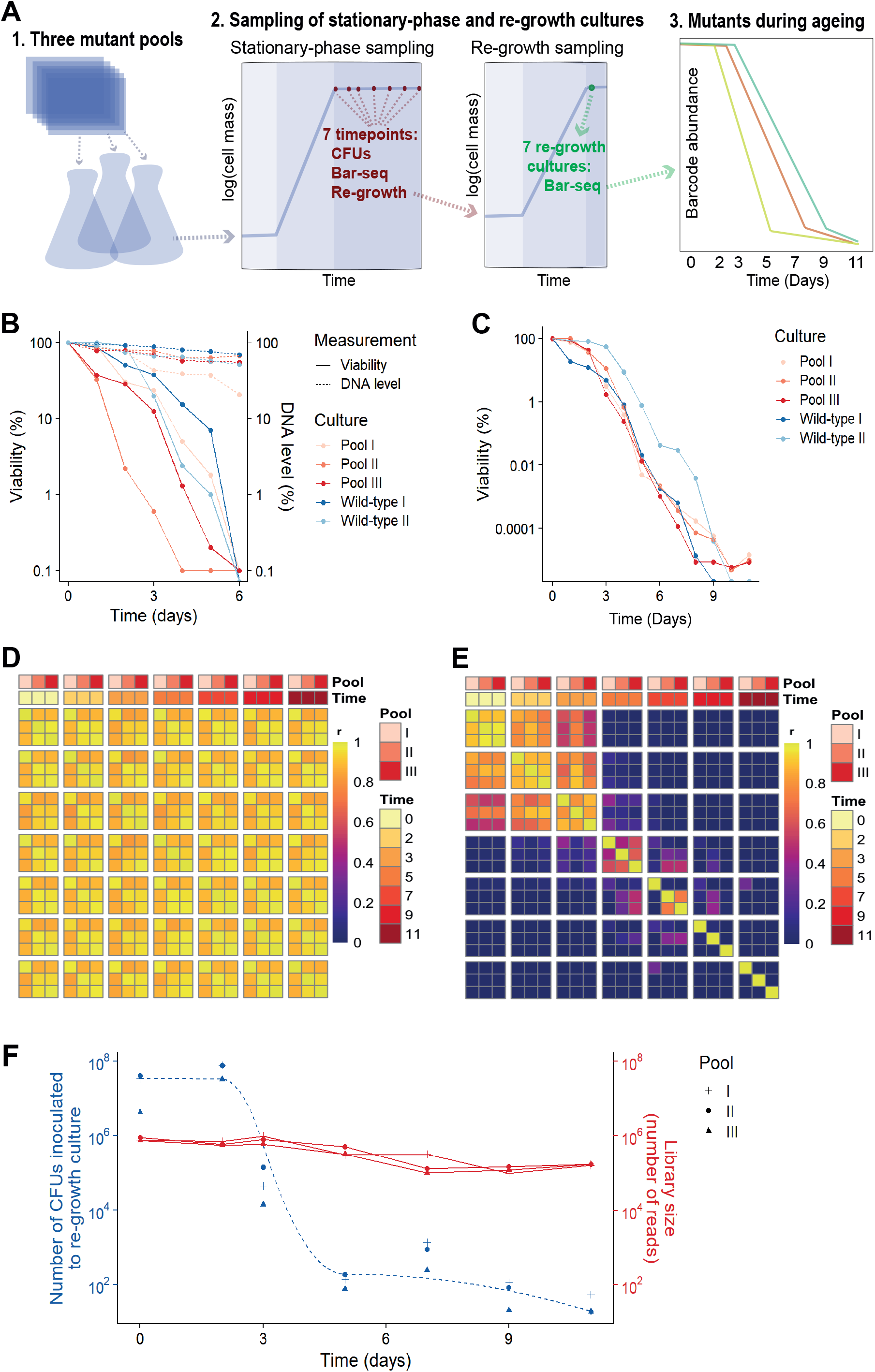
Bar-seq screen for CLS mutants. **A**. Scheme of Bar-seq CLS screen. 1. Three independent pools of prototroph deletion mutants were generated by re-suspending nine 384-colony plates in rich liquid medium as a pre-culture. 2. Pre-cultured cells were grown in fresh rich medium at 32°C for ∼2 days until saturation (100% cell viability), followed by sampling at indicated days to measure colony forming units (CFUs), to determine mutant abundance in aged cultures, and to inoculate fresh medium for determining mutant abundance after re-growth. 3. Selected samples were analysed by Bar-seq to identify long- and short-lived mutants. For the Bar-seq analyses, the following sample timepoints were used for all three repeats: Days 0, 2, 3, 5, 7, 9, 11. **B**. Experiment showing that DNA persists in dying cells. Pools of deletion mutants (three independent repeats) and wild-type control cells (two repeats) were grown in rich medium to stationary phase (Day 0), followed by measurements of cellular viability (CFU method) and DNA content (Qubit) as indicated. The data are relative to Day 0 which was set to 100%. **C**. CLS for the three independent pools of deletion mutant assays of the experiments used for the Bar-seq screen, along with CLS for two independent wild-type control cultures. The viability was determined using the CFU method and is indicated in log scale. Viability at the beginning of stationary phase (Day 0) was set to 100%, with viability of subsequent timepoints calculated relative to Day 0. **D**. Stationary-phase samples that were directly sequenced showed highly correlated barcode abundance across all timepoints and repeats (independent pools). Sample correlation between barcode counts from each pool across timepoints was calculated and plotted with the pheatmap package in R. **E**. Samples that were re-grown before sequencing showed substantially lower correlations in barcode abundance between timepoints (from Day 3) and even between repeats (from Day 5). Sample correlation between barcode counts is visualized as in D. **F**. Library size and number of live cells (CFUs) inoculated for each timepoint of the three re-growth experiments. From Day 3, the library size for a sample was greater than the number of cells inoculated for re-growth at that timepoint, leading to a sampling bottleneck that required scaling.

The ageing mutants that were directly sequenced before the re-growth showed highly correlated barcode abundance between all timepoints (Figure 1D). This result indicates that these samples do hardly capture differences in viability between mutants, consistent with DNA persisting in dead cells (Figure 1B). When re-growing the ageing mutant before sequencing, however, we did observe substantial ageing-related changes in relative abundance between mutants, reflecting different lifespans in different mutants (Figure 1E). Specifically, mutant abundances were highly correlated between Days 0 to 2, suggesting that most mutants remain viable at these timepoints, while these correlations began to fall apart from Day 3, suggesting that these samples become enriched for long-lived mutants. This result is consistent with a strong drop in viability of the stationary-phase pool around Day 3 (Figure 1C). These analyses show that mutants need to be re-grown before sampling by Bar-seq to restrict contribution from dead or non-proliferative cells.

### Late re-growth timepoints feature sampling bottleneck

After Day 5, mutant abundances composition in re-growth samples showed low correlations even between replicate pools of the same timepoint (Figure 1E). This poor correlation could reflect that mutant composition at these late timepoints is determined by stochastic sampling of few remaining mutants. To test this possibility, we used the CFU measurements of the stationary-phase cultures to estimate how many live cells were inoculated into the re-growth cultures at each timepoint (Figure 1F). This analysis showed that ∼100 or less live cells were present at the start of the re-growth cultures at Day 5 or later. Hence, inoculating re-growth cultures can introduce a substantial bottleneck at late timepoints, which must be accounted for to determine mutant abundance. In particular, when a re-growth culture is inoculated with a small number of progenitor cells, their clonal descendants from the same cell may be sequenced multiple times, resulting in overestimation of the statistical power. Therefore, where the library size for a sample was greater than the number of live cells inoculated for re-growth at that timepoint (Day 3 or later, Figure 1F), we scaled the read counts such that the library size equals the size of the bottleneck to ensure that each read represents on average one cell in the stationary-phase culture. Analogous conclusions have emerged from a recent study showing that barcode counts do not follow a negative binomial distribution in populations after strong selection bottlenecks, thus violating the statistical assumptions of RNA-seq algorithms typically employed for the analysis of count data [39]. We conclude that samples from late timepoints feature a technical bias, reflecting a sampling bottleneck which requires a special scaling procedure.

### Late stationary phase pools are biased by factors other than longevity

We considered which timepoints will maximise our ability to detect long- and short-lived mutants. The pools at the two last timepoints, Days 9 and 11, contained 29 mutants with an abundance of at least 1% of the read counts in one or more libraries. The results were stochastic, however, with the dominant mutants showing poor reproducibility between replicate pools at Days 9 and 11 (Figure 1E; Figure S2A). Notably, these 29 mutants typically decreased in abundance in early timepoints but then increased in abundance following the death of most other mutants (Figure S2B). The early decrease in abundance was statistically significant for 21 of these mutants (Figure S2C). Furthermore, the logFC between Day 3 and Day 0 for these 29 mutants was significantly lower than for all other mutants, revealing that pools at late timepoints were enriched for mutants classified as short-lived according to the earlier timepoints (Figure S2D). These results suggest that the persistence of these mutants at late timepoints reflects factors unrelated to longevity. For example, the nutrients released from dying mutants might be scavenged and provide a survival advantage to certain other mutants in a heterogeneous cell culture, a phenomenon that has been described in bacteria [40, 41], and recently in *S. pombe* cells during quiescence [42]. We conclude that samples from very late timepoints are also biased by biological phenomena that do not reflect longevity.

### Deletion mutants with altered chronological lifespans during stationary phase

Collectively, our analyses showed that mutants need to be re-grown before sampling by Bar-seq and that samples from the last timepoints can be biased through technical and biological effects which compromise the reliable detection of long-lived mutants. Hence, we limited our primary analysis to Day 0 (when 100% of cells were viable) and Day 3 (when ∼2% of cells were viable; Figure 1C). Our Bar-seq screen could detect 3061 mutants out of the 3206 decoded mutants (Table S2). For identifying long- and short-lived mutants, we analysed the normalised re-growth samples from Days 0 and 3 to estimate a fold change for each mutant. We used the following fold-change (FC) and false discovery rate (FDR) cut-offs for both long- and short-lived mutants: |log_2_(FC)| >log_2_(1.5) and FDR <0.05 (Figure 2A). This analysis identified 341 long-lived and 1246 short-lived mutants (Table S3).

**Figure 2:**
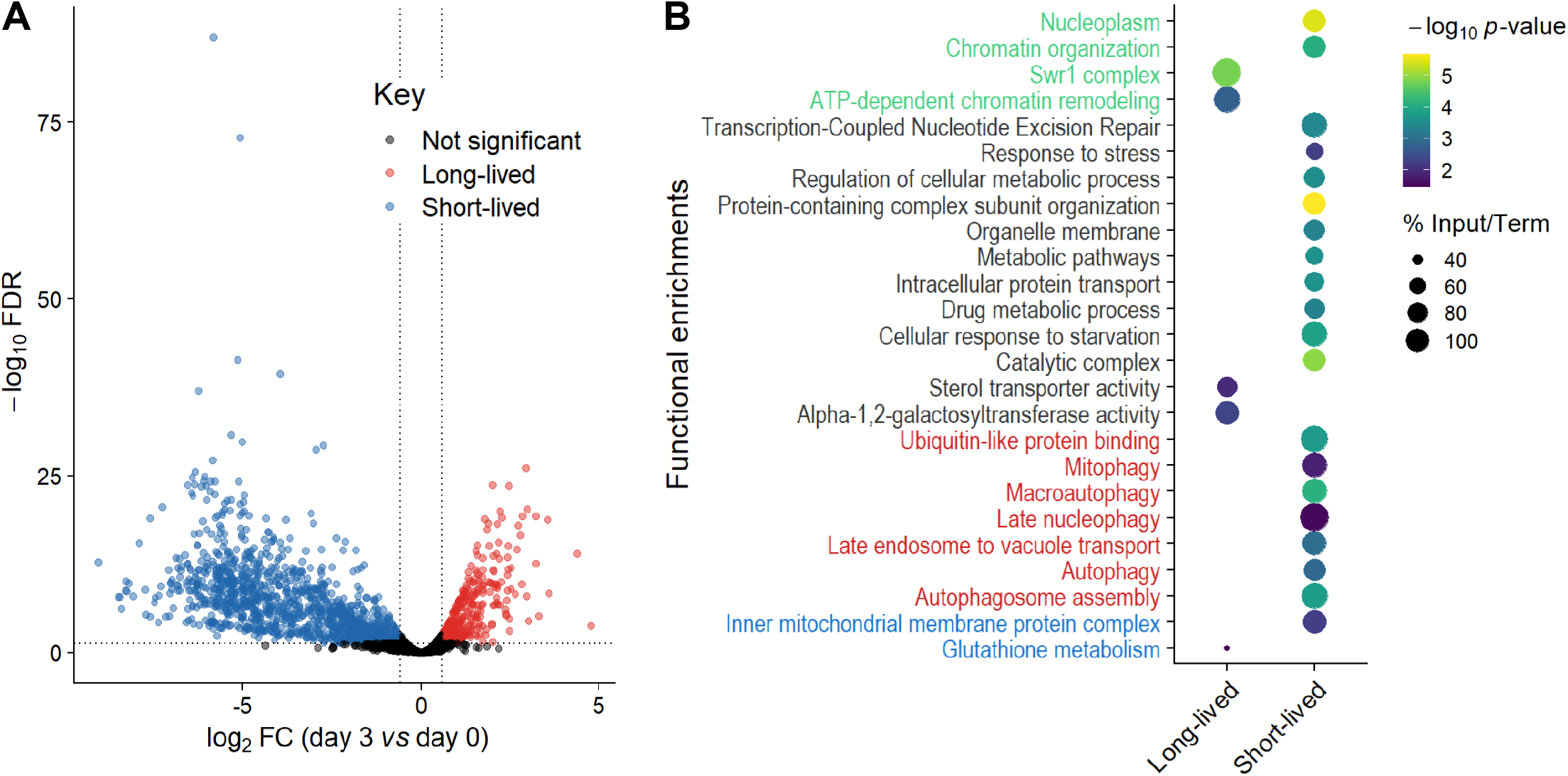
Long- and short-lived mutants and their functional enrichments. **A**. Volcano plot of mutant differences on Day 3 relative to Day 0 (log_2_ fold-change), based on Bar-seq of re-growth experiment, using DeSeq2 analysis of three independent repeats. Significance was determined using a fold-change (FC) cut-off of |log_2_(FC)| >log_2_(1.5) and a false discovery rate (FDR) cut-off of <0.05. **B**. Selected functional enrichments from Metascape [85] are shown for long- and short-lived mutants, including chromatin-related terms (green), autophagy-related terms (red), mitochondrial-related terms (blue) and other terms (black). The colour scale indicates significance expressed as -log_10_ p-values, and the size of the dots reflects the percentage of the input genes among all genes associated with the respective particular GO term.

We looked for functional enrichments among the genes which affect CLS. The short-lived mutants (reflecting genes that prolong lifespan) were enriched for several broad terms such as metabolic pathways, catalytic complex, chromatin organisation, intracellular protein transport and protein-containing complex subunit organisation (Figure 2B; Table S4). Such enrichments may reflect that gene deletions can be harmful for non-dividing cells by interfering with several different cellular processes, including those not directly related to ageing [19]. We also found enrichments for functions previously associated with stationary-phase survival, including cellular response to starvation, response to stress and regulation of cellular metabolic process (Figure 2B; Table S4). These enrichments may reflect the need for cells to respond to environmental changes and re-program their metabolism to maintain viability under nutrient-depleted conditions [43]. Another process critical for stationary-phase survival is autophagy, which allows recycling of damaged or surplus biomolecules and plays key roles in ageing and disease [44, 45]. In yeast, the vacuole is the site of autophagy and serves as a nutrient reservoir and signalling hub which integrates information from nutrient sensors [46, 47]. Accordingly, short-lived mutants were enriched for different terms related to the autophagy, including autophagosome formation and late endosome-to-vacuole transport (Figure 2B; Table S4). Selective processes of autophagy were also enriched, such as late nucleophagy and mitophagy, suggesting that recycling of nuclear and mitochondrial components is particularly important for stationary-phase survival. Indeed, late nucleophagy is a vital starvation response and associated with degenerative diseases [44, 45]; defective mitochondria can shorten the CLS [48], and inherited human diseases with mitophagy defects feature ageing pathologies such as neurodegeneration [49]. Short-lived mutants were also enriched for other mitochondrial terms (Figure 2B; Table S4), such as inner mitochondrial membrane, consistent with respiration being required for stationary-phase survival [50]. In humans, a decline in mitochondrial function is associated with ageing and degenerative diseases [51], with non-dividing brain cells being particularly sensitive to age-related mitochondrial impairments [52].

The long-lived mutants (reflecting genes that shorten lifespan) were also enriched in processes associated with respiration, such as glutathione metabolism (Figure 2B; Table S4). Glutathione is an antioxidant which detoxifies reactive oxygen species (a by-product of respiration) and a key determinant of redox signalling [53]. How impairment of glutathione metabolism could increase CLS is unclear, but reactive oxygen species, antioxidants and redox signalling play complex and nuanced roles in ageing [54]. Indeed, impairment of glutathione synthesis in budding yeast has different effects on CLS depending on nutritional status [55]. Furthermore, long-lived mutants were enriched for alpha-1,2-galactosyltransferase activity, raising the possibility that changes in glycosylation status play a role in ageing. In humans, the protein glycosylation status changes with age, which is especially relevant in non-proliferative tissues such as the nervous system [56, 57]. For example, alterations in protein glycosylation profiles, most notably β-Amyloid, is an early indicator of Alzheimer’s disease [58]. Another enrichment amongst the long-lived mutants was sterol transporter activity. Whilst it is unclear how impairment of sterol transport could increase CLS, sterols play important roles in metabolism and homeostasis [59] and have recently been shown to mediate the beneficial effects of dietary restriction in flies [60]. The long-lived mutants were also enriched in regulatory functions, such as ATP-dependent chromatin remodelling, a process carried out by evolutionarily conserved nucleosome remodelling factors which affect genome function, ageing and disease [61–63]. In particular, the Swr1 complex, or SRCAP in human, was highly enriched (Figure 2B; Table S4). SRCAP is a histone-exchange complex that deposits the histone variant H2A.Z at promoter regions, with broad roles for gene regulation [64, 65]. The enrichment of specific chromatin-related functions among the long-lived mutants suggests that chromatin regulators, such as the Swr1 complex, are involved in cellular ageing, possibly by modulating gene expression. Notably, Swr1 complex mutants are also long-lived in budding yeast, with the Swr1 complex being required for lifespan extension by dietary restriction [37]. These findings suggest that some chromatin regulators may participate in a conserved regulatory network that promotes ageing.

### Development of a robotics-based CFU assay

CLS in ageing experiments is traditionally measured by plating cells at different dilution factors and counting CFUs [21, 66]. However, measuring CLS using the traditional CFU method is both time- and resource-consuming and can lead to variable results. To circumvent these issues, we have developed a quantitative, automated CFU assay to facilitate medium- to high-throughput CLS studies. This new assay involves serial dilution of ageing cultures using a liquid-handling robot, followed by spotting droplets of the diluted cultures in quadruplicate on solid plates using a pinning robot (Figure 3A). This procedure results in colony patterns which reflect the proportions of viable cells in the corresponding cultures (Figure 3B). The assay is in essence a spot dilution approach, similar in concept to other approaches that capture differences in CFUs between cultures [36, 37]. Such an assay has the advantage that all dilution factors are spotted on a single agar plate, and multiple samples can be parallelised and analysed on the same plate. Hence, this new assay is much less resource- and time-demanding than the traditional CFU assay. For example, using the traditional assay to measure the lifespan of 24 ageing cultures at 10 timepoints (plating 3 dilution factors, with technical triplicates of each dilution factor) would require ∼2200 round agar plates, whilst the robotics-based assay could acquire the data using only 30 rectangular agar plates. Furthermore, the traditional CFU method can become experimentally unmanageable and intractable for studies containing more than ∼10 samples in parallel. We find that the ease at which our new assay can be implemented means that medium-scale ageing studies can now be readily and reliably conducted.

**Figure 3:**
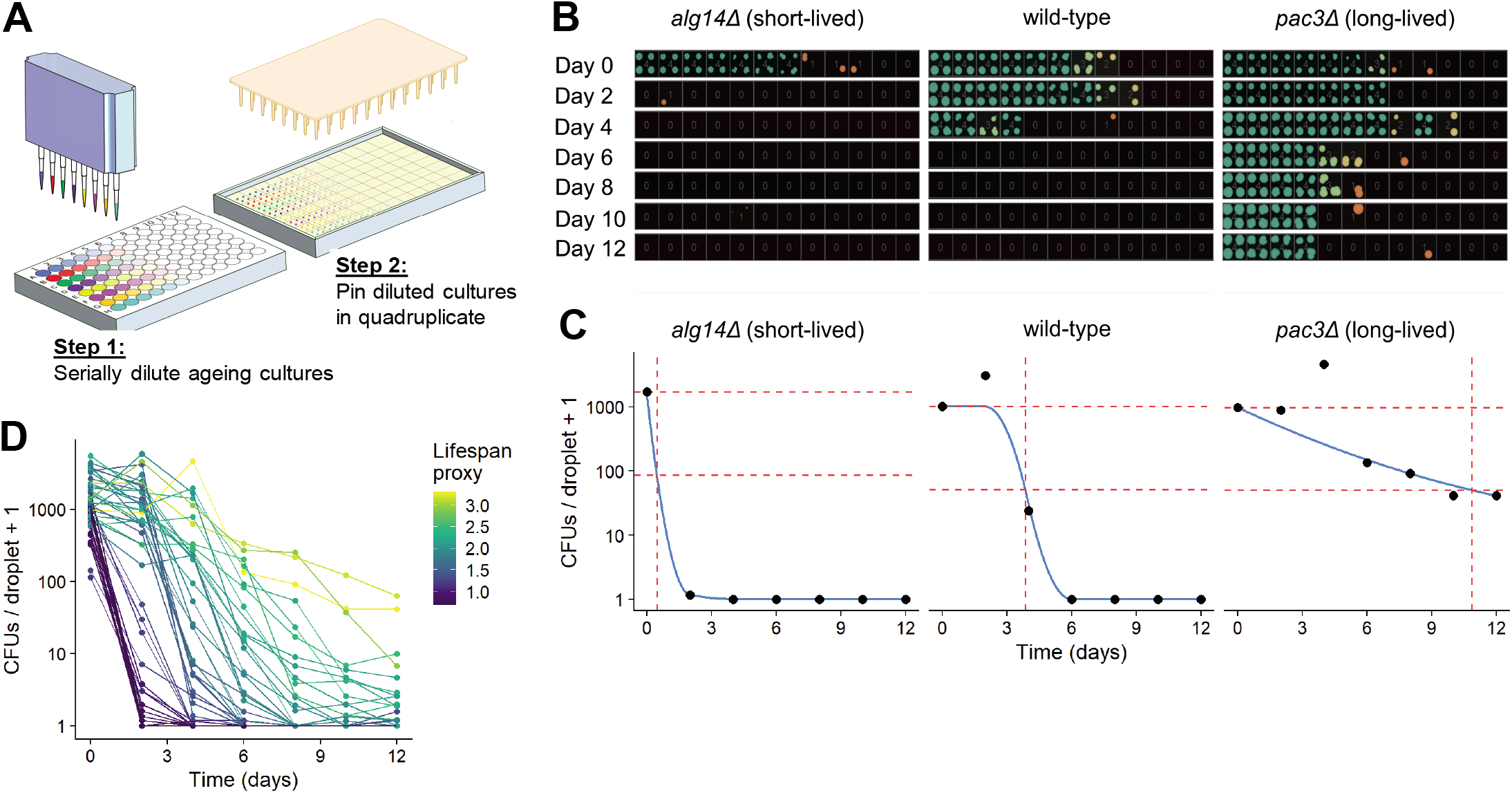
Development of robotics-based CFU assay. **A**. Scheme of high-throughput CFU protocol. Aliquots of ageing cultures are loaded into the first column of a 96-well plate (8 in parallel) and serially diluted 3-fold across the plate using a liquid handling robot or multichannel pipette. Droplets of diluted cultures are spotted onto solid agar in quadruplicate (384-well format) using a pinning robot. **B**. Agar plates are scanned after 2-4 days of growth, and images analysed using our R package, *DeadOrAlive*. Colony patterns for three strains (wild-type, short- and long-lived mutants predicted from Bar-seq screen) are shown for Days 0 to 12. Each position on the plate was scored for the presence or absence of colonies, and a vector extracted with digital information on the number of colonies for each dilution factor, sample, and timepoint. Colours from green through amber to red reflect decreasing colony numbers at each dilution factor (colony numbers are shown at the centre of each quadruplicate). **C**. Maximum likelihood estimates for number of CFUs/droplet plotted against time for the 3 samples in B. Blue lines show constrained smoothing spline fitted to each CLS curve. Red horizontal dashed lines represent numbers of CFUs/droplet at Day 0 (100% viability) and at 5% viability. Red vertical dashed line represents time at which 5% viability is reached according to the fitted values. The square root of this number was used as lifespan proxy for each sample. **D**. Maximum likelihood estimates of CFUs plotted against time for all 47 mutants validated from the Bar-seq screen, plus wild-type control. A CLS proxy was calculated for each sample (C), with each curve being coloured according to the proxy.

It can be conceptually and statistically challenging to analyse images of spot dilutions and quantitatively infer the number of CFUs in the ageing culture. Our variation of the spot-dilution assay solves this issue and provides quantitative estimates of CFUs for each sample. Each diluted sample is pinned multiple times and each position on the agar plate (in 384-well format) is scored for the presence or absence of a colony. Thus, a digital pattern representing the number of colonies at each dilution can be extracted for each sample and each timepoint (Figure 3B). To this end, we have developed an image analysis pipeline in R, based on the *gitter* package [67], to analyse plate images and extract colony patterns for each sample at each timepoint. The patterns can then be analysed using a statistical model to infer the number of CFUs via maximum likelihood estimation. This approach involves modelling the mean number of CFUs per culture droplet pinned onto the agar plate. Given that cultures are serially diluted prior to pinning, we assume that the mean number of cells per droplet exponentially decreases across the dilution factors. The number of cells pinned for each dilution factor can be modelled as Poisson distributed. Hence, the probability that a colony is present at each dilution is the sum of all probabilities for which at least 1 CFU has been pinned, and the probability that a colony is not present is the probability that no CFU has been pinned (Figure S3A). Given that each dilution factor has been pinned in quadruplicate, we can model the number of colonies present at each dilution factor as binomially distributed (Figure S3B).

To estimate the number of CFUs based on the observed patterns, a maximum-likelihood estimation is then performed to determine the number of CFUs per droplet of undiluted culture which is most likely to give rise to the observed patterns (Figure 3C). As with other maximum-likelihood estimators, this model is not robust to the presence of outliers; so we developed an algorithm that can identify and remove anomalous data points arising from errors such as plate contaminations or misclassifications by the image analysis. To estimate the CLS based on these CFUs and to facilitate comparison with other studies, we have also developed a proxy which describes the lifespan of a culture as a single value. To this end, we fit a constrained smoothing spline to the CFU data using the *cobs* package in R [68] (Figure 3C). Using this fit, we calculate the time taken for the culture viability to decrease to 5%. We use the square root of this number as the proxy value, as this proxy effectively captured differences in viability between long- and short-lived mutants (Figure 3D). All code to perform image analysis, maximum likelihood estimation and downstream analyses is available in our open-source R package, *DeadOrAlive* (https://github.com/JohnTownsend92/DeadOrAlive).

### Validation of robotics-based CFU assay against the traditional assay

In order to validate our new method, we measured CFUs for 6 strains with known differences in lifespan using both the traditional and robotics-based CFU method. Both methods recorded similar lifespan curves for each strain (Figure 4A), and the CFUs determined by the traditional method strongly correlated with the CFUs determined by the robotics-based assay across all timepoints (Figure 4B). Note, however, that the limit of detection was reached at earlier timepoints for the robotics-based method than the traditional method, meaning that the high-throughput method cannot capture differences in CFUs for cultures of very low cell viabilities (Figure 4A). We conclude that the robotics-based method can reliably estimate CFUs and, therefore, facilitate the measurement of CLS for large numbers of samples.

**Figure 4:**
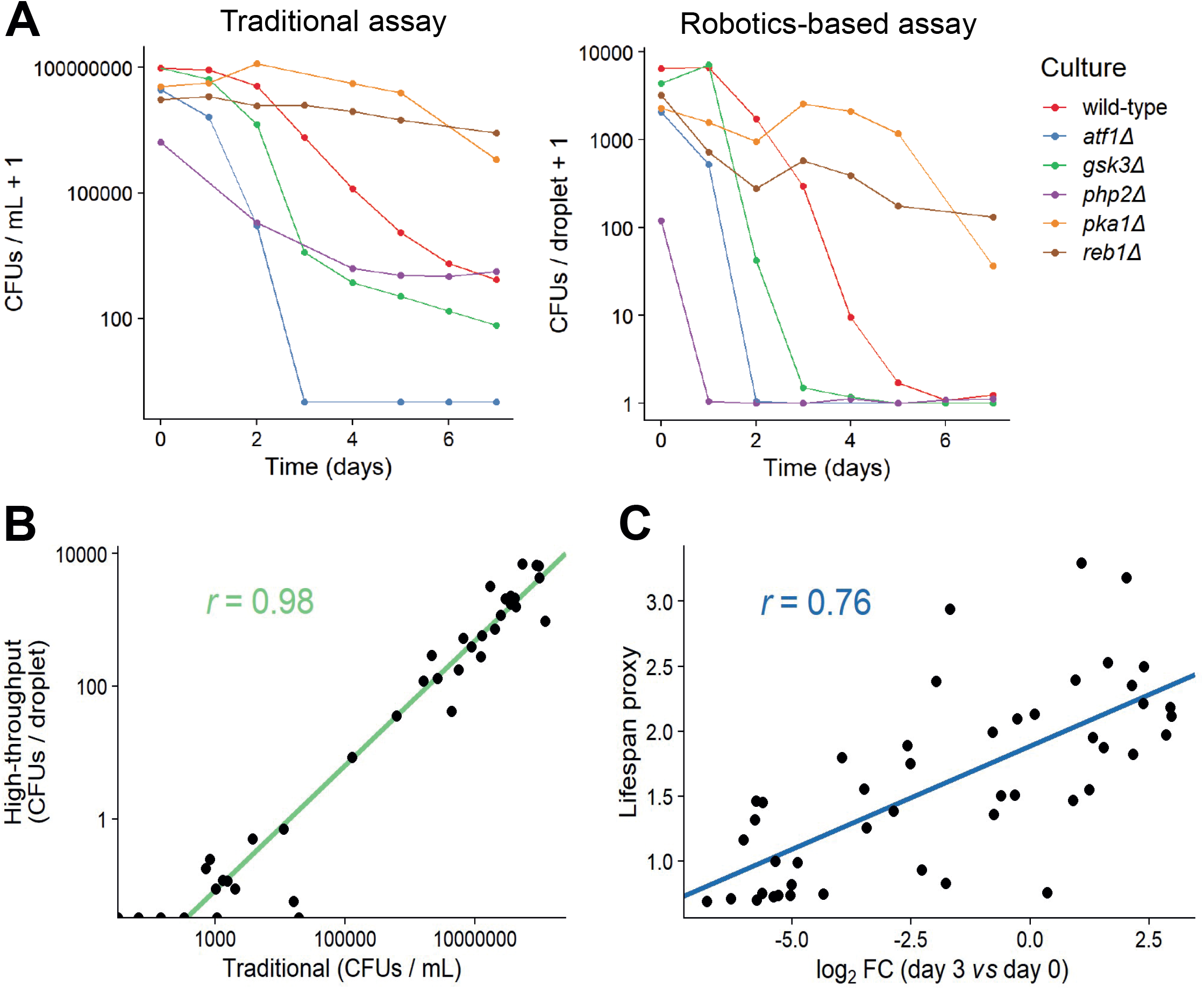
Comparison of traditional and high-thoughput lifespan assays. **A**. Comparison of traditional and robotics-based CFU assays. Lifespan curves for 6 mutants with different lifespans were measured in parallel using both the traditional (left) and robotics-based (right) CFU assays. Both methods capture differences in lifespan between short-lived (*atf1Δ, gsk3Δ, php2Δ*) and long-lived (*pka1Δ, reb1Δ*) mutants. **B**. CFUs measured for various cultures (different mutants grown under different conditions) at different timepoints, using both the traditional and robotics-based method. Scatter plot shows the agreement between the two methods. Green: linear regression of log-transformed CFU values (traditional *vs* robotics-based), along with Pearson correlation coefficient. **C**. Scatter plot showing the agreement of CLS estimates between Bar-seq and the robotics-based CFU assay for the 47 validated mutants. The log_2_ fold change (FC) of barcode abundance (Day 3 relative to Day 0) is plotted against the lifespan proxy based on maximum-likelihood estimates from the robotics-based CFU method. Blue: linear regression between log_2_ FC and lifespan proxy, along with Pearson correlation coefficient.

### Validation of selected mutants from Bar-seq screen using robotics-based CFU assay

We then exploited the new CFU assay to validate the CLS data from the Bar-seq screen. Figure 3B shows the pattern of colonies produced by two strains featuring strong antagonistic effects of CLS in the Bar-seq data: a new short-lived mutant (*alg14*) and a new long-lived mutant (*pac3*), along with wild-type control cells. Figure 3C shows maximum likelihood estimates for the number of CFUs for these three strains based on the observed colony patterns and the fitted constrained smoothing spline to calculate the time taken for cell viability to decrease to 5%. This analysis confirmed that the two mutants showed the CLS effects expected based on the Bar-seq data (Figure 3C).

We then applied the robotics-based CFU assay to validate the CLS of 47 mutants that showed a range of lifespans in the Bar-seq screen, including two known short-lived mutants (*sdh1* and *coq5*) and three known long-lived mutants (*tco89, pyp1* and *git3*), along with wild-type control cells (Table S5). To facilitate comparison between the two datasets, we applied our proxy to reduce the dimensionality and summarise the lifespan based on the shape of the survival curve (Figure 3C,D). Using this proxy, we compared the results of the validation to the original Bar-seq screen, revealing substantial overall agreement between the two methods (Figure 4C). This finding is reassuring given that the two methods employ distinct experimental and analytical approaches. We conclude that the Bar-seq screen was successful in uncovering mutants with altered CLS.

### New ageing-associated genes identified in Bar-seq screen

We compared the ageing-associated genes identified in the Bar-seq screen with known ageing-related genes (Table S3). Overall, 166 of our 1587 hits have previously been associated with fission yeast phenotype ontologies indicating altered CLS, including ‘increased viability in stationary phase/upon nitrogen starvation’ and ‘loss of viability in stationary phase/G0/upon nitrogen starvation/nutrient depletion/glucose starvation’ [14, 69]. For example, 55 and 21 hits have been identified as ageing mutants in screens for altered CLS during quiescence [19] or for mutants resistant to TORC1 inhibitors [20], respectively. Moreover, 266 hits are listed in the GenAge database as ageing-related genes in different organisms [70]. Although these overlaps are substantial, our screen also uncovered an excess of genes not previously implicated in ageing. Notably, 51 hits included ‘priority unstudied genes’, a set of ∼140 genes that are conserved from fission yeast to human but have not been directly studied in any organism [71]. This result raises the intriguing possibility that many of these unstudied genes actually play roles in ageing-related processes, as has been speculated [71]. Of the 47 independently validated genes (Fig. 3E), 33 mutants have not previously been associated with ageing, including 10 ‘pro-ageing’ genes and 23 ‘anti-ageing’ genes (Table S5). Characterization of these genes might enlighten unknown yet conserved processes of cellular ageing. Interestingly, among the novel ‘pro-ageing’ proteins, Jac1, SPCC1494.08c, Cyp4 and Rpl1102 have human orthologs implicated in disease [14]. These orthologs include HSCB, a co-chaperone involved in iron-sulphur cluster formation, which is associated with increased susceptibility to ataxia [72]; FAM102A, which has a putative role in estrogen action [73] and is implicated in a type of glaucoma [74]; PPIB, a endoplasmic-reticulum isomerase involved in collagen biosynthesis and linked to osteogenesis imperfecta [75, 76]; and RPL11, a ribosomal protein associated with Diamond-Blackfan anaemia [77].

## Conclusion

We decoded barcodes for 3206 mutants of the latest *S. pombe* deletion library (ver 5.0), most of which for both barcodes, enabling Bar-seq screening of a substantially increased number of genes. We established an improved experimental and analytical pipeline to facilitate Bar-seq assays in general, and CLS screens in particular, addressing technical and statistical issues raised by sampling at late timepoints and by the re-growth protocol needed because DNA persists in dead cells. We identified 341 long-lived and 1246 short-lived deletion mutants that point to a large number of new ageing-associated genes, including 51 conserved but entirely uncharacterized genes. We also developed a robotics-based CFU assay and analysis pipeline, facilitating medium- to high-throughput CLS studies of batch cultures. We used this assay to validate the lifespan of 47 mutants identified in the Bar-seq screen, revealing good agreement despite substantial differences in biological context (ageing in pool *vs* batch cultures) and experimental approaches (relative barcode abundance in regrowth cultures *vs* CFUs). Our validation uncovered 33 new genes not previously associated with cellular ageing. This study provides potent systematic approaches and new genes to study cellular ageing.

## MATERIALS AND METHODS

### Pooling and growth of deletion strains

Prototroph and auxotroph strain pools of the latest *S. pombe* gene-deletion library (ver. 5.0; Bioneer, South Korea) were generated as described [19]. The prototroph library was combined in a single pool for CLS screening. The auxotroph library, used only for the barcode decoding, was divided into 9 separate pools for each plate (in 384-well format) in order to maximise the PCR amplification and decoding of mutants. For all mutant pools, sample collection and storage were processed in the same manner. Pool aliquots of 500 µL, stored at -80°C at a final concentration of 20% (v/v) glycerol, were thawed on ice, cells were re-suspended in 250 mL YES medium [78] at a density of ∼1.0 OD_600nm_ in 500 mL conical flasks, with pre-cultures grown at 25°C for ∼14-16 hours without shaking. Cells were washed and re-suspended to 0.2 OD_600nm_ in the required volume of YES. Cultures were grown to stationary phase at 32°C and 170 rpm for 2 days, unless stated otherwise, at which point cultures were considered to be 100% viable. Once stationary phase was reached, culture viability was determined as described [21]. In parallel, re-growth cultures were inoculated and grown until stationary phase. Aliquots of 2 mL were washed as before and the pellets stored at -80°C until required for DNA extraction for both ageing cultures and re-growth cultures.

### Library preparation and sequencing

Genomic DNA was extracted using the MasterPure Yeast DNA Purification Kit (Epicentre, UK). During the extraction protocol, a lysis step was introduced as follows: cells were lysed twice with mechanical beating using glass beads (0.5 mm diameter, Stratech Scientific, UK) in a FastPrep-24 Instrument (MP Biomedicals, UK) and incubated for 1 hour at 65°C. DNA was purified and quantified using the QIAquick PCR purification kit (Qiagen, UK) and Qubit (ThermoFisher Scientific, Rochford, UK), respectively, following the manufacturer’s instructions.

For barcode decoding, DNA from two independent aliquots of the auxotroph deletion-strain pool were used. Purified DNA (25 ng/µL) in 100 µL of nuclease-free water (Qiagen, UK) was broken down to ∼400 bp with 7 cycles of 30 seconds shearing and 30 seconds rest using the Diagenode Bioruptor® instrument (ATG Scientific, UK). Barcodes were treated separately by end-repair using the NEBNext® End Repair Module (NEB, UK), linker ligation using the NEBNext® Quick Ligation Module (NEB, UK) and amplified with Phusion® High-Fidelity DNA polymerase (NEB, UK) using its dedicated master mix (NEB, UK) and custom-designed primers. These primers consisted of linkers required for extracting genomic sequences and barcode-specific sequences. Linker oligo sequences were: 5’ – TTCAGACGTGTGCTCTTCCGATCTNNNNNNNNNNCAGGCTACTCCGCTTAAGGGAC-3’ (linker 1, Invitrogen, UK) and 5’-GTCCCTTAAGCGGAGTAGCCTG/3AmMO/-3’ (linker 2, DNA IDT, UK). Both UpTag and DnTag forward primer sequences were complementary to linker 1. Reverse UpTag primer 5’ – <u>CACGACGCTCTTCCGATCT</u>**<u>AGTA</u>**NNNNGGGGACGAGGCAAGCTAAGATATC-3’ (Invitrogen, UK) and reverse DnTag 5’ – <u>CACGACGCTCTTCCGATCT</u>**AGTA**NNNNCGCCATCCAGTGTCGAAAAGTATC-3’ (Invitrogen, UK) primer sequences comprised part of the Illumina adaptor sequence (underlined), four constant bases (‘AGTA’), four random bases (‘Ns’) acting as unique molecular identifiers (UMIs), and the U2/D2 UpTag- and DnTag-specific sequences. DNA was amplified with 15 cycles of 10 seconds at 98°C, 45 seconds at 65°C and 30 seconds at 72°C. DNA was diluted ten-fold and used as template for the second round of PCR where Illumina adaptors for sequencing were added using the NEBNext® Multiplex Oligos Illumina dual index kit (NEB, UK) with 10 cycles of 10 seconds at 98°C, 45 seconds at 65°C and 30 seconds at 72°C. Size selection for fragments of approximately 450-550 bp was performed using AMPure® XP beads (NEB, UK). Briefly, AMPure beads were incubated at room temperature for 30 mins. Size selection was performed at 1.2x (beads volume/sample volume) in a total volume of 100 µL, followed by incubation at room temperature for 5 mins before placing on a magnetic stand to separate the beads and discard the supernatant. The beads with the DNA were then washed twice with 200 µL freshly prepared 80% EtOH, air dried for approximately 5 mins, and DNA eluted in 25 µL nuclease-free H_2_O (Qiagen). When required, a further 1x (beads volume/sample volume) was performed to remove any leftover primer dimers. Libraries were quantified with Qubit and run on the BioAnalyzer (Agilent, UK). Sequencing was performed on an Illumina MiSeq Instrument (Illumina, US) with 168 cycles using paired-end reads of 75 bp each generating approximately 30 million reads.

For CLS screening, DNA was extracted (using MasterPure Yeast DNA Purification Kit; Epicentre, UK) from stationary phase and re-growth cultures for selected timepoints (Days 0, 2, 3, 5, 7, 9 and 11). Starting with 250 ng of total DNA, UpTag and DnTag barcodes were separately amplified with Phusion® High-Fidelity DNA polymerase (NEB, UK) using custom-designed primers at a concentration of 100 nM each in a total volume of 50 µL, with 6 cycles of 10 seconds at 98°C, 30 seconds at 60°C and 30 seconds at 72°C. Oligo sequences of UpTag and DnTag (Invitrogen, UK) were: 5’ – <u>TTCAGACGTGTGCTCTTCCGATCT</u>GTCANNNNCGCTCCCGCCTTACTTCGCATTTAAA-3’ and 5’-<u>CACGACGCTCTTCCGATCT</u>AGTANNNNGGGGACGAGGCAAGCTAAGATATC-3’, and 5’-<u>CACGACGCTCTTCCGATCT</u>AGTANNNNCGCCATCCAGTGTCGAAAAGTATC-3’ and 5’-<u>TTCAGACGTGTGCTCTTCCGATCT</u>GTCANNNNTTGCGTTGCGTAGGGGGGATTTAAA-3’, respectively. These sequences were custom-designed and differed from previously described barcode sequencing methods [19, 33] by containing part of the Illumina adaptor sequence (underlined), four constant bases (‘GTCA’ or ‘AGTA’) introduced to easily identify the start of the reads, four random bases ‘Ns’ added to act as Unique Molecular Identifiers (UMIs), U1/U2 and D2/D1 UpTag- and DnTag-specific sequences. Products were purified using the MinElute® PCR Purification Kit (Qiagen, Germany) and eluted in 10 µL dH_2_O. All 10 µL of the purified product was used as template for the second PCR in a total volume of 25 µL with 17 cycles of 10 seconds at 98°C, 30 seconds at 65°C and 30 seconds at 72°C using the NEBNext® Multiplex Oligos kit (NEB, UK). The expected library size was ∼200-250bp. To select for this range, we removed fragments <150 bp using 1x AMPure® XP beads (NEB, UK). DNA quantification and quality control was performed using a BioAnalyser Instrument (Agilent Technologies, US). Libraries were pooled at a total concentration of 4 nM, and PhiX sequencing control v3 (Illumina, US) to increase the library complexity was added at a concentration of 5%. Libraries were sequenced on an Illumina MiSeq Instrument with 168 cycles using paired-end reads of 75 bp each and generating approximately 30 million reads.

### Decoding of Deletion Library Barcodes

Figure S4A provides a scheme of the steps taken to decode the barcodes of the ver. 5.0 deletion library (Bioneer). Reads from each of the 9 pool plates were combined and analysed collectively. The paired-end reads (R1 and R2) were analysed separately. UpTag and DnTag R1 reads were mapped to the respective UpTag or DnTag barcode flank sequences, U1/U2 (5’-CAAGCTAAGATATC-3’ and 5’-TTTAAATGCGAAGTAA-3’) and D2/D1 (5’-AGTGTCGAAAAGTATC-3’ and 5’ – TTTAAAATCCCCCCTA-3’). The barcode sequence was extracted from between these flanks. UpTag and DnTag reads containing some primer sequence as part of the genomic DNA were removed by mapping the R2 reads to the primer sequences, and the genomic DNA was extracted as the R2 sequence minus the primer sequence. Mapping to flanking/primer sequences and identification of barcodes/genomic DNA was performed using an in-house Python script, *Barcount* (https://github.com/Bahler-Lab/barcount). To ensure that genomic DNA fragments were genuine, we used the FASTQX-Toolkit [79] to filter sequences against the UpTag/DnTag (U1/D2) primer sequences, thus removing possible primer contaminated genomic sequences. Genomic DNA reads were mapped to the *S. pombe* reference genome using Bowtie2 [80]. Next, we used BEDtools [81] to identify the nearest upstream/downstream gene to the mapped region for the UpTag/DnTag respectively, taking into account the directionality of genes. We discarded reads where a barcode could not be extracted from the R1 read or the R2 read could not be uniquely mapped to a gene. Figure S4B shows the read loss following the different steps of these analyses.

In order to match barcodes to genes with high confidence, we identified barcode-gene pairs which appeared in reads with high frequency. This was performed separately for UpTag and DnTag barcodes (Figure S5). In order to account for possible indels or base mutations known to arise within synthetic barcodes sequences [36], pairwise Levenshtein distance was calculated between all barcodes, and barcodes were assembled into clusters where they differed by no more than 3 mutations. A consensus barcode was defined for each cluster as the average barcode sequence of that cluster. A consensus barcode was automatically assigned to a gene if the following 3 criteria were met: 1) there were at least 10 reads where a consensus barcode mapped to a gene; 2) at least 80% of all reads containing a consensus barcode mapped to a gene; and 3) at least 80% of all reads mapped to a gene associated with a consensus barcode. A subset of the automatically assigned barcode-gene pairs were manually inspected using an in-house genome browser, where the number of reads for UpTags and DnTags was plotted with respect to genome position. This browser was also used to inspect and manually assign cases where automatic assignment was not possible, such as overlapping genes. Code for the creation of consensus barcodes, the assignment of barcode-gene pairs, and the in-house genome browser are available in the *BarSeqTools* R package (https://github.com/Catalina37/Barcount_BarSeqTools_Pipelines/tree/master/BarSeqTools).

### Application of Bar-seq to Identify Long- and Short-lived Mutants

Paired-end reads were assembled using PEAR [82] and filtered for PCR duplicates using BEDTools [83]. Barcodes for UpTags and DnTags were extracted by mapping reads to the respective UpTag/DnTag flanking sequences using *Barcount*. A code-example of how *Barcount* for UpTags was run is as follows: barcount --fastq UpTag.fastq --rmdup --flanking_left CAAGCTAAGATATC -- flanking_right TTTAAATGCGAAGTAA --max_distance_flanks 1 --max_distance_barcode 3 -- barcode_table UpTagReference.csv --debug --verbose --save_extracted_barcodes --out UpTag.filter.fastq. A code-example of running *Barcount* for the DnTags is as follows: barcount -- fastq DnTag.fastq --rmdup --flanking_left AGTGTCGAAAAGTATC --flanking_right TTTAAAATCCCCCCTA --max_distance_flanks 1 --max_distance_barcode 3 --barcode_table DnTagReference.csv --debug --verbose --save_extracted_barcodes --out DnTag.filter.fastq.

Barcodes were matched to genes according to the lookup table compiled, and a total read count for each gene was created by adding up counts for UpTag and DnTag. The number of CFUs present in the ageing cultures at each timepoint was used to estimate the size of the bottleneck introduced by inoculating re-growth cultures (i.e. how many live cells were used to inoculate the re-growth culture). If the library size was greater than the bottleneck size, read counts were scaled prior to differential fitness analysis to ensure that the library size equalled the bottleneck size, ensuring that the scaled read counts represented the number of live cells present in the stationary phase culture at each timepoint. Differential fitness analysis based on barcode frequency in the re-growth cultures was performed using edgeR (version 3.24.3) [84]. Time was considered as a categorical variable, and the pool was included as a term in the model in order to account for differences in barcode abundance between pools. Read counts were modelled using a negative binomial generalised linear model with likelihood ratio testing being used to determine *p*-values for differences in barcode abundances between timepoints. For determination of long- and short-lived mutants, timepoints 0 and 3 were analysed using a fold-change (FC) cut-off of |log_2_(FC)| > log_2_(1.5) and false discovery rate (FDR) cut-off of FDR < 0.05. Enrichment analyses of long- and short-lived gene deletion lists were performed with Metascape [85] and AnGeLi [86]. In both cases, the 3061 genes whose effect on lifespan we could measure in the Bar-seq screen were used as the background for enrichment tests.

### Development of a robotics-based CFU assay

As described in Results, we developed a novel assay to measure CFUs from batch cell cultures which can be largely automated by robotics. We loaded 150 µL aliquots of ageing culture into the first column of a 96-well plate (8 cultures in parallel per plate). The rest of the plate was loaded with 100 µL of YES. By taking 50 µL of the ageing culture from the first column, cultures were serially diluted 3-fold across the plate, ensuring each dilution factor was well mixed before proceeding to the next. This was performed using an Integra Assist automated multichannel pipette (Integra Biosciences Ltd). Droplets of serially diluted ageing cultures were immediately dispensed onto YES agar in quadruplicate (384-well format) using a Singer RoToR HDA pinning robot (Singer Instruments). For this, long-pin 96-density pads were used, making sure that the source plate was revisited before each pin onto agar. Plates were incubated for 2-4 days at 32°C until patterns of colonies appeared. Images of agar plates were acquired with pyphe-scan [87] using an Epson V700 scanner in transmission mode. We provide an R package, *DeadOrAlive*, to analyse images of plates and quantify the number of CFUs in the ageing culture based on the colony patterns observed.

### Validation of robotics-based CFU assay against the traditional assay

In order to validate the CFU assay, we measured the lifespan of 6 different strains grown in YES using both the traditional and high-throughput methods in parallel. Cultures were grown to stationary phase at 32°C and 170 rpm for 2 days, at which point cultures were considered to be 100% viable. Once stationary phase was reached, culture viability was determined as described previously [21] and using the high-throughput method described above. For wild-type, the *972 h-* strain was used, *gsk3::natMX6 h-* was generated in a previous study [88], whilst *reb1::natMX6 h-, php2::natMX6 h-, atf1::natMX6 h-* and *pka1::kanMX4 h-* were generated as described previously [89].

### Validation of selected mutants from Bar-seq screen using robotics-based CFU assay

Excluding the wild-type control (*972 h*^*-*^), we selected 47 mutants in total with varying lifespans from the Bar-seq screen to independently validate the lifespans using the robotics-based CFU assay. Apart from the wild-type cells, which were independently grown, all mutant strains were manually selected from fresh prototrophic cell colonies, grown on YES plates, re-streaked onto new YES plates, and grown at 32°C for 2 days. Colonies were used to set individual pre-cultures grown in parallel in 20 mL YES overnight at 32°C and 170 rpm. Cultures of 20 mL YES at 0.002 OD_600nm_ were prepared from the corresponding pre-cultures and grown to saturation at 32°C with 170 rpm shaking. Once cultures reached saturation, the first timepoint (Day 0, 100% cell viability) was collected and processed using the robotics-based CFU assay as described earlier.

## ACKNOWLEDGMENTS

We thank Mimoza Hoti for advice and help with some experiments and Pawan Dhami for help with sequencing (Genomics and Genome Engineering Facility funded by the Cancer Research UK-UCL Centre). This work was supported by the Biotechnology and Biological Sciences Research Council [grant number BB/M009513/1], by a Boehringer Ingelheim Fonds PhD fellowship to StJ.T., and by a Welcome Trust Senior Investigator Award to J.B. (Grant No. 095598/Z/11/Z). The Francis Crick Institute receives its core funding from Cancer Research UK (FC001134), the UK Medical Research Council (FC001134) and the Wellcome Trust (FC001134).

## References

1. Partridge L, and Gems D (2002). Mechanisms of aging: public or private? Nat Rev Genet. 3(3): 165–175. doi: 10.1038/nrg753.

2. López-Otín C, Blasco MA, Partridge L, Serrano M, and Kroemer G (2013). The hallmarks of aging. Cell 153:1194.

3. Niccoli T, and Partridge L (2012). Ageing as a Risk Factor for Disease. Curr Biol. 22(17): R741–R752. doi: 10.1016/J.CUB.2012.07.024.

4. Kenyon C, Chang J, Gensch E, Rudner A, and Tabtiang R (1993). A C. elegans mutant that lives twice as long as wild type. Nature. 366(6454): 461–464. doi: 10.1038/366461a0.

5. Kapahi P, Chen D, Rogers AN, Katewa SD, Li PWL, Thomas EL, and Kockel L (2010). With TOR, less is more: A key role for the conserved nutrient-sensing TOR pathway in aging. Cell Metab. 11:453–465.

6. Katewa SD, and Kapahi P (2010). Dietary restriction and aging, 2009. Aging Cell. 9(2): 105–112. doi: 10.1111/j.1474-9726.2010.00552.x.

7. Longo VD, and Finch CE (2003). Evolutionary Medicine: From Dwarf Model Systems to Healthy Centenarians? Science. 299(5611): 1342–1346. doi: 10.1126/science.1077991.

8. De Magalhães JP, and Tacutu R (2015). Integrative Genomics of Aging. In: Handb. Biol. Aging Eighth Ed. pp 263–285.

9. Lucanic M et al. (2017). Impact of genetic background and experimental reproducibility on identifying chemical compounds with robust longevity effects. Nat Commun. 8(1): 1–13. doi: 10.1038/ncomms14256.

10. Phillips P, Lithgow GJ, and Driscoll M (2017). A long journey to reproducible results. Nature 548:387–388.

11. Rallis C, and Bähler J (2016). Cell-based screens and phenomics with fission yeast. Crit Rev Biochem Mol Biol. 51(2): 86–95. doi: 10.3109/10409238.2015.1103205.

12. Fruhmann G, Seynnaeve D, Zheng J, Ven K, Molenberghs S, Wilms T, Liu B, Winderickx J, and Franssens V (2017). Yeast buddies helping to unravel the complexity of neurodegenerative disorders. Mech Ageing Dev. 161(Pt B): 288–305. doi: 10.1016/j.mad.2016.05.002.

13. Kaeberlein M (2010). Lessons on longevity from budding yeast. Nature. 464(7288): 513–519. doi: 10.1038/nature08981.

14. Lock A, Rutherford K, Harris MA, Hayles J, Oliver SG, Bähler J, and Wood V (2019). PomBase 2018: User-driven reimplementation of the fission yeast database provides rapid and intuitive access to diverse, interconnected information. Nucleic Acids Res. 47(D1): D821–D827. doi: 10.1093/nar/gky961.

15. Pancaldi V, Schubert F, and Bähler J (2010). Meta-analysis of genome regulation and expression variability across hundreds of environmental and genetic perturbations in fission yeast. Mol Biosyst. 6(3): 543–552. doi: 10.1039/b913876p.

16. Ohtsuka H, Shimasaki T, and Aiba H (2020). Genes affecting the extension of chronological lifespan in Schizosaccharomyces pombe (fission yeast). Mol Microbiol. doi: 10.1111/mmi.14627.

17. Ellis DA, Mustonen V, Rodríguez-López M, Rallis C, Malecki M, Jeffares DC, and Bähler J (2018). Uncovering Natural Longevity Alleles from Intercrossed Pools of Aging Fission Yeast Cells. Genetics. 210(2): 733–744. doi: 10.1534/genetics.118.301262.

18. Masuda F, Ishii M, Mori A, Uehara L, Yanagida M, Takeda K, and Saitoh S (2016). Glucose restriction induces transient G2 cell cycle arrest extending cellular chronological lifespan. Sci Rep. 6(1): 19629. doi: 10.1038/srep19629.

19. Sideri T, Rallis C, Bitton DA, Lages BM, Suo F, Rodríguez-López M, Du L-L, and Bähler J (2015). Parallel Profiling of Fission Yeast Deletion Mutants for Proliferation and for Lifespan During Long-Term Quiescence. G3 Genes Genomes Genetics. 5(1): 145–155. doi: 10.1534/g3.114.014415.

20. Rallis C, López-Maury L, Georgescu T, Pancaldi V, and Bähler J (2014). Systematic screen for mutants resistant to TORC1 inhibition in fission yeast reveals genes involved in cellular ageing and growth. Biol Open. 3(2): 161–71. doi: 10.1242/bio.20147245.

21. Rallis C, Codlin S, and Bähler J (2013). TORC1 signaling inhibition by rapamycin and caffeine affect lifespan, global gene expression, and cell proliferation of fission yeast. Aging Cell. 12(4): 563–73. doi: 10.1111/acel.12080.

22. Aranda-Anzaldo A (2012). The post-mitotic state in neurons correlates with a stable nuclear higher-order structure. Commun Integr Biol. 5(2): 134–9. doi: 10.4161/cib.18761.

23. Zuin A, Carmona M, Morales-Ivorra I, Gabrielli N, Vivancos AP, Ayté J, and Hidalgo E (2010). Lifespan extension by calorie restriction relies on the Sty1 MAP kinase stress pathway. EMBO J. 29(5): 981–991. doi: 10.1038/emboj.2009.407.

24. Roux AE, Leroux A, Alaamery MA, Hoffman CS, Chartrand P, Ferbeyre G, and Rokeach LA (2009). Pro-Aging Effects of Glucose Signaling through a G Protein-Coupled Glucose Receptor in Fission Yeast. PLoS Genet. 5(3). doi: 10.1371/JOURNAL.PGEN.1000408.

25. Roux AE, Quissac A, Chartrand P, Ferbeyre G, and Rokeach LA (2006). Regulation of chronological aging in Schizosaccharomyces pombe by the protein kinases Pka1 and Sck2. Aging Cell. 5(4): 345–357. doi: 10.1111/j.1474-9726.2006.00225.x.

26. Gulli J, Cook E, Kroll E, Rosebrock A, Caudy A, and Rosenzweig F (2019). Diverse conditions support near-zero growth in yeast: Implications for the study of cell lifespan. Microb. Cell 6:397–413.

27. Barré BP, Hallin J, Yue JX, Persson K, Mikhalev E, Irizar A, Holt S, Thompson D, Molin M, Warringer J, and Liti G (2020). Intragenic repeat expansion in the cell wall protein gene HPF1 controls yeast chronological aging. Genome Res. 30(5): 697–710. doi: 10.1101/gr.253351.119.

28. Belak ZR, Harkness T, and Eskiw CH (2018). A rapid, high-throughput method for determining chronological lifespan in budding yeast. J Biol Methods. 5(4). doi: 10.14440/jbm.2018.272.

29. Khawaja A-A-W, Belak ZR, Eskiw CH, and Harkness TAA (2021). High-Throughput Rapid Yeast Chronological Lifespan Assay. Methods Mol Biol Clifton NJ. 2196: 229–233. doi: 10.1007/978-1-0716-0868-5_18.

30. Powers RW, Kaeberlein M, Caldwell SD, Kennedy BK, Fields S, and Fields S (2006). Extension of chronological life span in yeast by decreased TOR pathway signaling. Genes Dev. 20(2): 174–84. doi: 10.1101/gad.1381406.

31. Avelar-Rivas JA, Munguía-Figueroa M, Juárez-Reyes A, Garay E, Campos SE, Shoresh N, and DeLuna A (2020). An Optimized Competitive-Aging Method Reveals Gene-Drug Interactions Underlying the Chronological Lifespan of Saccharomyces cerevisiae. Front Genet. 11: 468. doi: 10.3389/fgene.2020.00468.

32. Matecic M, Smith DL, Pan J, Maqani X, and Bekiranov N (2010). A Microarray-Based Genetic Screen for Yeast Chronological Aging Factors. PLoS Genet. 6(4): 1000921. doi: 10.1371/journal.pgen.1000921.

33. Han TX, Xu X-Y, Zhang M-J, Peng X, and Du L-L (2010). Global fitness profiling of fission yeast deletion strains by barcode sequencing. Genome Biol. 11(6): R60. doi: 10.1186/gb-2010-11-6-r60.

34. Kim D-U et al. (2010). Analysis of a genome-wide set of gene deletions in the fission yeast MSchizosaccharomyces pombe. Nat Biotechnol. 28(6): 617–623. doi: 10.1038/nbt.1628.

35. Smith DL, Maharrey CH, Carey CR, White RA, Hartman JL, Hartman JL, and IV (2016). Gene-nutrient interaction markedly influences yeast chronological lifespan. Exp Gerontol. 86: 113–123. doi: 10.1016/j.exger.2016.04.012.

36. Lie S, Banks P, Lawless C, Lydall D, and Petersen J (2018). The contribution of non-essential Schizosaccharomyces pombe genes to fitness in response to altered nutrient supply and target of rapamycin activity. Open Biol. 8(5): 180015. doi: 10.1098/rsob.180015.

37. Garay E, Campos SE, González de la Cruz J, Gaspar AP, Jinich A, and DeLuna A (2014). High-Resolution Profiling of Stationary-Phase Survival Reveals Yeast Longevity Factors and Their Genetic Interactions. PLoS Genet. 10(2): e1004168. doi: 10.1371/journal.pgen.1004168.

38. Fabrizio P, Hoon S, Shamalnasab M, Galbani A, Wei M, Giaever G, Nislow C, and Longo VD (2010). Genome-Wide Screen in Saccharomyces cerevisiae Identifies Vacuolar Protein Sorting, Autophagy, Biosynthetic, and tRNA Methylation Genes Involved in Life Span Regulation. PLoS Genet. 6(7): e1001024. doi: 10.1371/journal.pgen.1001024.

39. Akimov Y, Bulanova D, Timonen S, Wennerberg K, and Aittokallio T (2020). Improved detection of differentially represented DNA barcodes for high‐throughput clonal phenomics. Mol Syst Biol. 16(3). doi: 10.15252/msb.20199195.

40. Bacun-Druzina V, Cagalj Z, and Gjuracic K (2007). The growth advantage in stationary-phase (GASP) phenomenon in mixed cultures of enterobacteria. FEMS Microbiol Lett. 266(1): 119–27. doi: 10.1111/j.1574-6968.2006.00515.x.

41. Finkel SE (2006). Long-term survival during stationary phase: Evolution and the GASP phenotype. Nat. Rev. Microbiol. 4:113–120.

42. Makarenko R, Denis C, Francesconi S, Gangloff S, and Arcangioli B (2020). Nitrogen starvation reveals the mitotic potential of mutants in the S/MAPK pathways. Nat Commun. 11(1): 1–13. doi: 10.1038/s41467-020-15880-y.

43. Zhang N, and Cao L (2017). Starvation signals in yeast are integrated to coordinate metabolic reprogramming and stress response to ensure longevity. Curr Genet. 63(5): 839–843. doi: 10.1007/s00294-017-0697-4.

44. Wong SQ, Kumar A V., Mills J, and Lapierre LR (2020). Autophagy in aging and longevity. Hum Genet. 139(3): 277–290. doi: 10.1007/s00439-019-02031-7.

45. Papandreou ME, and Tavernarakis N (2019). Nucleophagy: from homeostasis to disease. Cell Death Differ. 26(4): 630–639. doi: 10.1038/s41418-018-0266-5.

46. Aufschnaiter A, and Büttner S (2019). The vacuolar shapes of ageing: From function to morphology. Biochim Biophys Acta -Mol Cell Res. 1866(5): 957–970. doi: 10.1016/j.bbamcr.2019.02.011.

47. Carmona-Gutierrez D, Hughes AL, Madeo F, and Ruckenstuhl C (2016). The crucial impact of lysosomes in aging and longevity. Ageing Res Rev. 32: 2–12. doi: 10.1016/j.arr.2016.04.009.

48. Plummer JD, and Johnson JE (2019). Extension of Cellular Lifespan by Methionine Restriction Involves Alterations in Central Carbon Metabolism and Is Mitophagy-Dependent. Front Cell Dev Biol. 7. doi: 10.3389/fcell.2019.00301.

49. Bakula D, and Scheibye-Knudsen M (2020). MitophAging: Mitophagy in Aging and Disease. Front Cell Dev Biol. 8. doi: 10.3389/fcell.2020.00239.

50. Ocampo A, Liu J, Schroeder EA, Shadel GS, and Barrientos A (2012). Mitochondrial Respiratory Thresholds Regulate Yeast Chronological Life Span and its Extension by Caloric Restriction. Cell Metab. 16(1): 55–67. doi: 10.1016/j.cmet.2012.05.013.

51. Sun N, Youle RJ, and Finkel T (2016). The Mitochondrial Basis of Aging. Mol Cell. 61(5): 654–666. doi: 10.1016/j.molcel.2016.01.028.

52. Grimm A, and Eckert A (2017). Brain aging and neurodegeneration: from a mitochondrial point of view. J Neurochem. 143(4): 418–431. doi: https://doi.org/10.1111/jnc.14037.

53. Lu SC (2013). Glutathione synthesis. Biochim Biophys Acta. 1830(5): 3143–3153. doi: 10.1016/j.bbagen.2012.09.008.

54. Hekimi S, Lapointe J, and Wen Y (2011). Taking a “good” look at free radicals in the aging process. Trends Cell Biol. 21(10): 569–576. doi: 10.1016/j.tcb.2011.06.008.

55. Tello‐Padilla MF, Perez‐Gonzalez AY, Canizal‐García M, González‐Hernández JC, Cortes‐Rojo C, Olivares‐Marin IK, and Madrigal‐Perez LA (2018). Glutathione levels influence chronological life span of Saccharomyces cerevisiae in a glucose-dependent manner. Yeast. 35(5): 387–396. doi: https://doi.org/10.1002/yea.3302.

56. Ando S (2014). Glycoconjugate Changes in Aging and Age-Related Diseases. In: Yu RK, Schengrund C-L, editors Glycobiol. Nerv. Syst. Springer, New York, NY; pp 415–447.

57. Sato Y, and Endo T (2010). Alteration of brain glycoproteins during aging. Geriatr Gerontol Int. 10(1): S32–S40. doi: https://doi.org/10.1111/j.1447-0594.2010.00602.x.

58. Regan P, McClean PL, Smyth T, and Doherty M (2019). Early Stage Glycosylation Biomarkers in Alzheimer’s Disease. Medicines. 6(3): 92. doi: 10.3390/medicines6030092.

59. Wollam J, and Antebi A (2011). Sterol Regulation of Metabolism, Homeostasis, and Development. Annu Rev Biochem. 80(1): 885–916. doi: 10.1146/annurev-biochem-081308-165917.

60. Zanco B, Mirth CK, Sgrò CM, and Piper MD (2021). A dietary sterol trade-off determines lifespan responses to dietary restriction in Drosophila melanogaster females. eLife. 10: e62335. doi: 10.7554/eLife.62335.

61. Sewda A, Agopian AJ, Goldmuntz E, Hakonarson H, Morrow BE, Taylor D, and Mitchell LE (2019). Gene-based genome-wide association studies and meta-analyses of conotruncal heart defects. PLoS ONE. 14(7). doi: 10.1371/journal.pone.0219926.

62. Purohit JS, and Chaturvedi MM (2016). Chromatin and aging. In: Top. Biomed. Gerontol. Springer Singapore; pp 205–241.

63. Liu BK.H Yip R, and Zhou Z (2012). Chromatin Remodeling, DNA Damage Repair and Aging. Curr Genomics. 13(7): 533–547. doi: 10.2174/138920212803251373.

64. Clément‐Ziza M, Marsellach FX, Codlin S, Papadakis MA, Reinhardt S, Rodríguez‐López M, Martin S, Marguerat S, Schmidt A, Lee E, Workman CT, Bähler J, and Beyer A (2014). Natural genetic variation impacts expression levels of coding, non‐coding, and antisense transcripts in fission yeast. Mol Syst Biol. 10(11): 764. doi: 10.15252/msb.20145123.

65. Wong MM, Cox LK, and Chrivia JC (2007). The chromatin remodeling protein, SRCAP, is critical for deposition of the histone variant H2A.Z at promoters. J Biol Chem. 282(36): 26132–26139. doi: 10.1074/jbc.M703418200.

66. Chen B-R, and Runge KW (2009). A New Schizosaccharomyces pombe Chronological Lifespan Assay Reveals that Caloric Restriction Promotes Efficient Cell Cycle Exit and Extends Longevity. Exp Gerontol. 44(8): 493–502. doi: 10.1016/j.exger.2009.04.004.

67. Wagih O, and Parts L (2014). Gitter: A robust and accurate method for quantification of colony sizes from plate images. G3 Genes Genomes Genet. 4(3): 547–552. doi: 10.1534/g3.113.009431.

68. Ng P, and Maechler M (2007). A fast and efficient implementation of qualitatively constrained quantile smoothing splines. In: Stat. Model. Sage Publications India Pvt. Ltd,B-42, Panchsheel Enclave, New Delhi; pp 315–328.

69. Harris MA, Lock A, Bähler J, Oliver SG, and Wood V (2013). FYPO: the fission yeast phenotype ontology. Bioinformatics. 29(13): 1671–1678. doi: 10.1093/bioinformatics/btt266.

70. Tacutu R, Thornton D, Johnson E, Budovsky A, Barardo Di, Craig T, DIana E, Lehmann G, Toren D, Wang J, Fraifeld VE, and De Magalhães JP (2018). Human Ageing Genomic Resources: New and updated databases. Nucleic Acids Res. 46(D1): D1083–D1090. doi: 10.1093/nar/gkx1042.

71. Wood V, Lock A, Harris MA, Rutherford K, Bähler J, and Oliver SG (2019). Hidden in plain sight: what remains to be discovered in the eukaryotic proteome? Open Biol. 9(2): 180241. doi: 10.1098/rsob.180241.

72. Sun G, Gargus JJ, Ta DT, and Vickery LE (2003). Identification of a novel candidate gene in the iron-sulfur pathway implicated in ataxia-susceptibility: Human gene encoding HscB, a J-type co-chaperone. J Hum Genet. 48(8): 415–419. doi: 10.1007/s10038-003-0048-9.

73. Bateman A (2019). UniProt: A worldwide hub of protein knowledge. Nucleic Acids Res. 47(D1): D506–D515. doi: 10.1093/nar/gky1049.

74. Nongpiur ME, Cheng CY, Duvesh R, Vijayan S, Baskaran M, Khor CC, Allen J, Kavitha S, Venkatesh R, Goh D, Husain R, Boey PY, Quek D, Ho CL, Wong TT, Perera S, Wong TY, Krishnadas SR, Sundaresan P, Aung T, and Vithana EN (2018). Evaluation of Primary Angle-Closure Glaucoma Susceptibility Loci in Patients with Early Stages of Angle-Closure Disease. Ophthalmology. 125(5): 664–670. doi: 10.1016/j.ophtha.2017.11.016.

75. Rappaport N, Twik M, Plaschkes I, Nudel R, Iny Stein T, Levitt J, Gershoni M, Morrey CP, Safran M, and Lancet D (2017). MalaCards: an amalgamated human disease compendium with diverse clinical and genetic annotation and structured search. Nucleic Acids Res. 45(D1): D877– D887. doi: 10.1093/nar/gkw1012.

76. van Dijk FS, Nesbitt IM, Zwikstra EH, Nikkels PGJ, Piersma SR, Fratantoni SA, Jimenez CR, Huizer M, Morsman AC, Cobben JM, van Roij MHH, Elting MW, Verbeke JIML, Wijnaendts LCD, Shaw NJ, Högler W, McKeown C, Sistermans EA, Dalton A, Meijers-Heijboer H, and Pals G (2009). PPIB Mutations Cause Severe Osteogenesis Imperfecta. Am J Hum Genet. 85(4): 521–527. doi: 10.1016/j.ajhg.2009.09.001.

77. Chakraborty A, Uechi T, Nakajima Y, Gazda HT, O’Donohue MF, Gleizes PE, and Kenmochi N (2018). Cross talk between TP53 and c-Myc in the pathophysiology of Diamond-Blackfan anemia: Evidence from RPL11-deficient in vivo and in vitro models. Biochem Biophys Res Commun. 495(2): 1839–1845. doi: 10.1016/j.bbrc.2017.12.019.

78. Moreno S, Klar A, and Nurse P (1991). Molecular genetic analysis of fission yeast Schizosaccharomyces pombe. Methods Enzymol. 194(C): 795–823. doi: 10.1016/0076-6879(91)94059-L.

79. Gordon, A., & Hannon GJ (2010). FASTX-Toolkit. http://hannonlab.cshl.edu/fastx_toolkit/index.html..

80. Langmead B, and Salzberg SL (2012). Fast gapped-read alignment with Bowtie 2. Nat Methods. 9(4): 357–359. doi: 10.1038/nmeth.1923.

81. Quinlan AR, and Hall IM (2010). BEDTools: A flexible suite of utilities for comparing genomic features. Bioinformatics. 26(6): 841–842. doi: 10.1093/bioinformatics/btq033.

82. Zhang J, Kobert K, Flouri T, and Stamatakis A (2014). PEAR: A fast and accurate Illumina Paired-End reAd mergeR. Bioinformatics. 30(5): 614–620. doi: 10.1093/bioinformatics/btt593.

83. Bushnell B (JGI) (2014). BBMap short read aligner, and other bioinformatic tools.

84. Robinson MD, Mccarthy DJ, and Smyth GK (2010). edgeR: a Bioconductor package for differential expression analysis of digital gene expression data. Bioinforma Appl NOTE. 26(1): 139–140. doi: 10.1093/bioinformatics/btp616.

85. Zhou Y, Zhou B, Pache L, Chang M, Khodabakhshi AH, Tanaseichuk O, Benner C, and Chanda SK (2019). Metascape provides a biologist-oriented resource for the analysis of systems-level datasets. Nat Commun. 10(1). doi: 10.1038/s41467-019-09234-6.

86. Bitton DA, Schubert F, Dey S, Okoniewski M, Smith GC, Khadayate S, Pancaldi V, Wood V, and Bähler J (2015). AnGeLi: A tool for the analysis of gene lists from fission yeast. Front Genet. 6(NOV): 330. doi: 10.3389/fgene.2015.00330.

87. Kamrad S, Rodríguez-López M, Cotobal C, Correia-Melo C, Ralser M, and Bähler J (2020). Pyphe, a python toolbox for assessing microbial growth and cell viability in high-throughput colony screens. eLife. 9: e55160. doi: 10.7554/eLife.55160.

88. Rallis C, Townsend S, and Bähler J (2017). Genetic interactions and functional analyses of the fission yeast gsk3 and amk2 single and double mutants defective in TORC1-dependent processes. Sci Rep. 7(1): 1–11. doi: 10.1038/srep44257.

89. Bähler J, Wu JQ, Longtine MS, Shah NG, McKenzie A, Steever AB, Wach A, Philippsen P, and Pringle JR (1998). Heterologous modules for efficient and versatile PCR-based gene targeting in Schizosaccharomyces pombe. Yeast. 14(10): 943–951. doi: 10.1002/(SICI)1097-0061(199807)14:10<943::AID-YEA292>3.0.CO;2-Y.

