## Supplementary material for "Barcode Sequencing and a High-throughput Assay for Chronological Lifespan Uncover Ageing-associated Genes in Fission Yeast": Figures S1 to S5

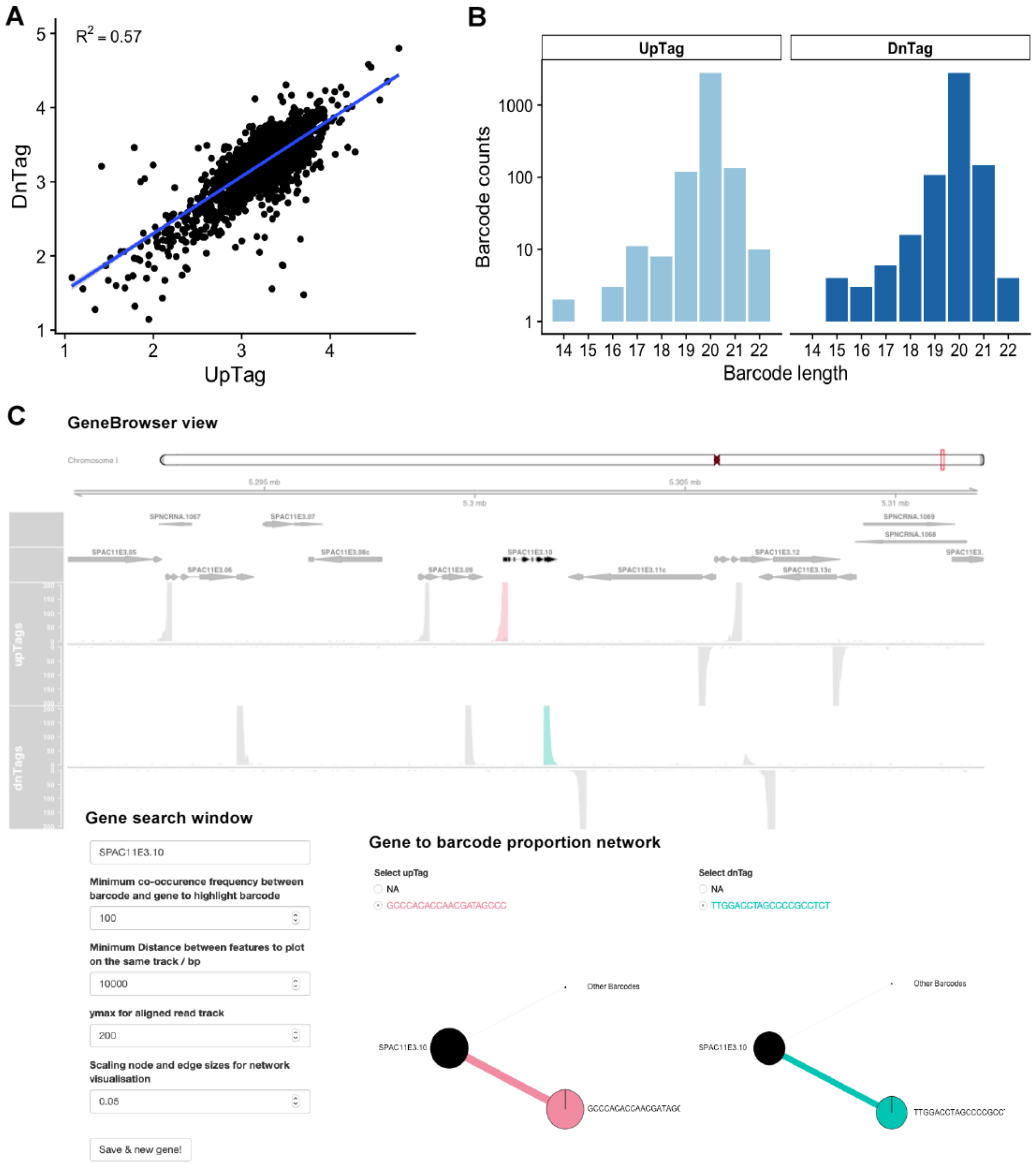


**Figure S1: Barcode decoding of *S. pombe* deletion-mutant library ver. 5.0.**
**A.** Correlation plot between UpTag and DnTag gene-barcode pair frequencies. Barcode counts (number of sequenced barcodes) are provided as $\log_{10}$ values. The correlation coefficient between the UpTag and DnTag counts was calculated using a linear model (p-adjusted <2.6E-16).
**B.** Counts of different sequence lengths of decoded UpTag and DnTag barcodes are shown as $\log_{10}$.
**C.** *GeneBrowser* view of a gene associated with both barcodes. The browser allows visualisation of the query gene and its available annotations. The tracks: protein-coding genes (grey), deleted/query gene (black), UpTag reads (cyan) and DnTag reads (pink). The directionality and position of barcodes relative to the query gene can be used for manual gene curation. The Gene search window (bottom left) contains the parameters that can be adjusted, while the Gene to barcode proportion network (bottom right) displays the percentage of a gene that maps to the aligned barcode(s). In this example, there is only one barcode which maps to the query gene with a proportion of 100% as shown by the pie charts.

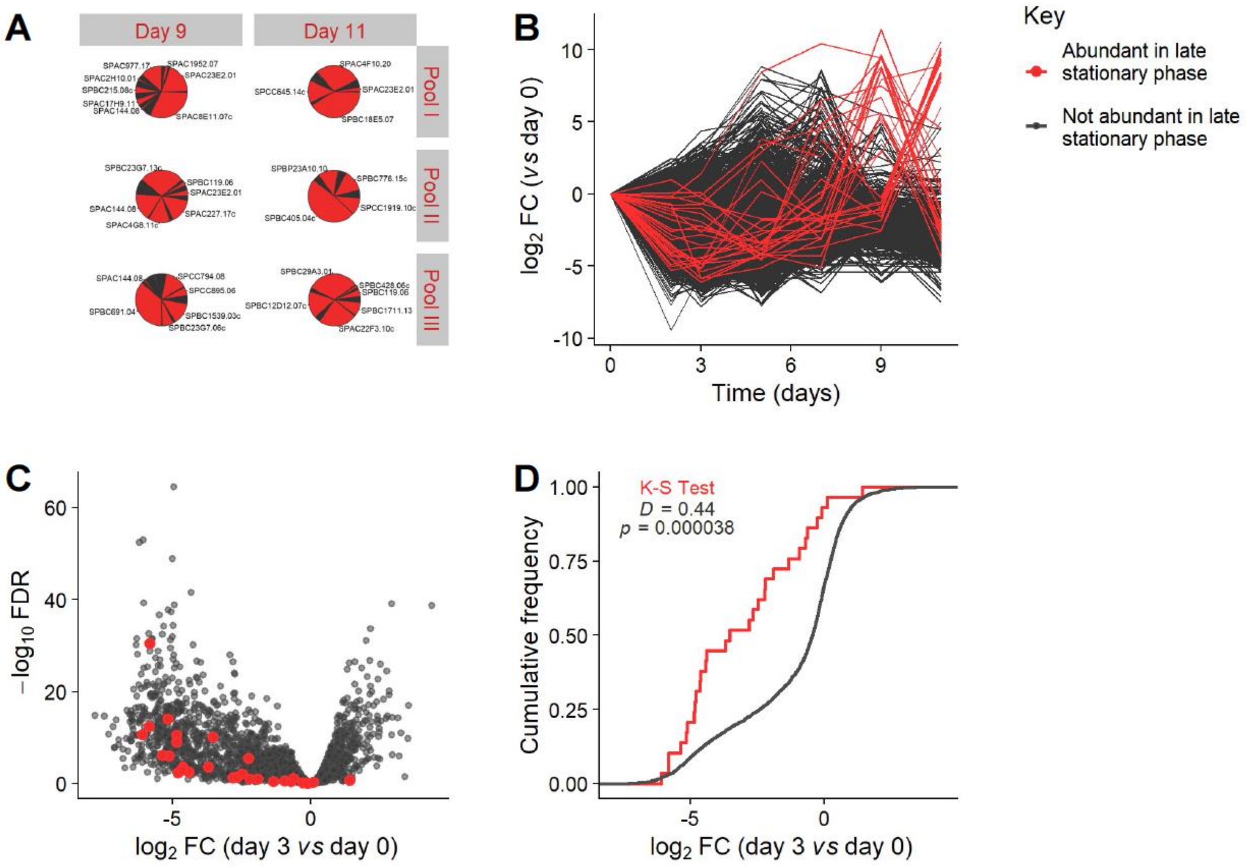


**Figure S2: Late stationary phase pools are dominated by short-lived mutants.**
**A.** Pie charts showing barcode abundances in re-growth pools in late stationary phase (Days 9 and 11). 29 mutants were defined as highly abundant in these pools (mutants which contributed >1% of the reads for at least one of the re-growth libraries at Days 9 or 11). These mutants are shown in red, while all other mutants are shown in dark grey. In each library, the vast majority of reads map to between 4 and 8 mutants.
**B.** Line plot where $log_2$ fold change (FC) of barcode abundances (each Day *vs* Day 0) is plotted against time for each mutant. Mutants dominating the late pools typically decrease earlier in stationary phase before increasing in late stationary phase.
**C.** Volcano plot of $log_2$ FC in early stationary phase (Day 3 *vs* Day 0). Barcode abundances for 21 of the 29 mutants (red) which dominate late stationary phase pools are significantly lower at Day 3 than at Day 0 ($log_2$ (FC) <-$log_2$ (1.5); FDR <0.05).
**D.** Cumulative frequency plot for distribution of $log_2$ FC (Day 3 *vs* Day 0). Mutants which dominate late stationary phase pools have a significantly lower $log_2$ FC than all other mutants (Kolmogorov-Smirnov test, $D$ = 0.44, $p$ = 0.00004). This indicates that late stationary phase pools are significantly enriched for mutants which are classified as short-lived according to the analysis of earlier timepoints.

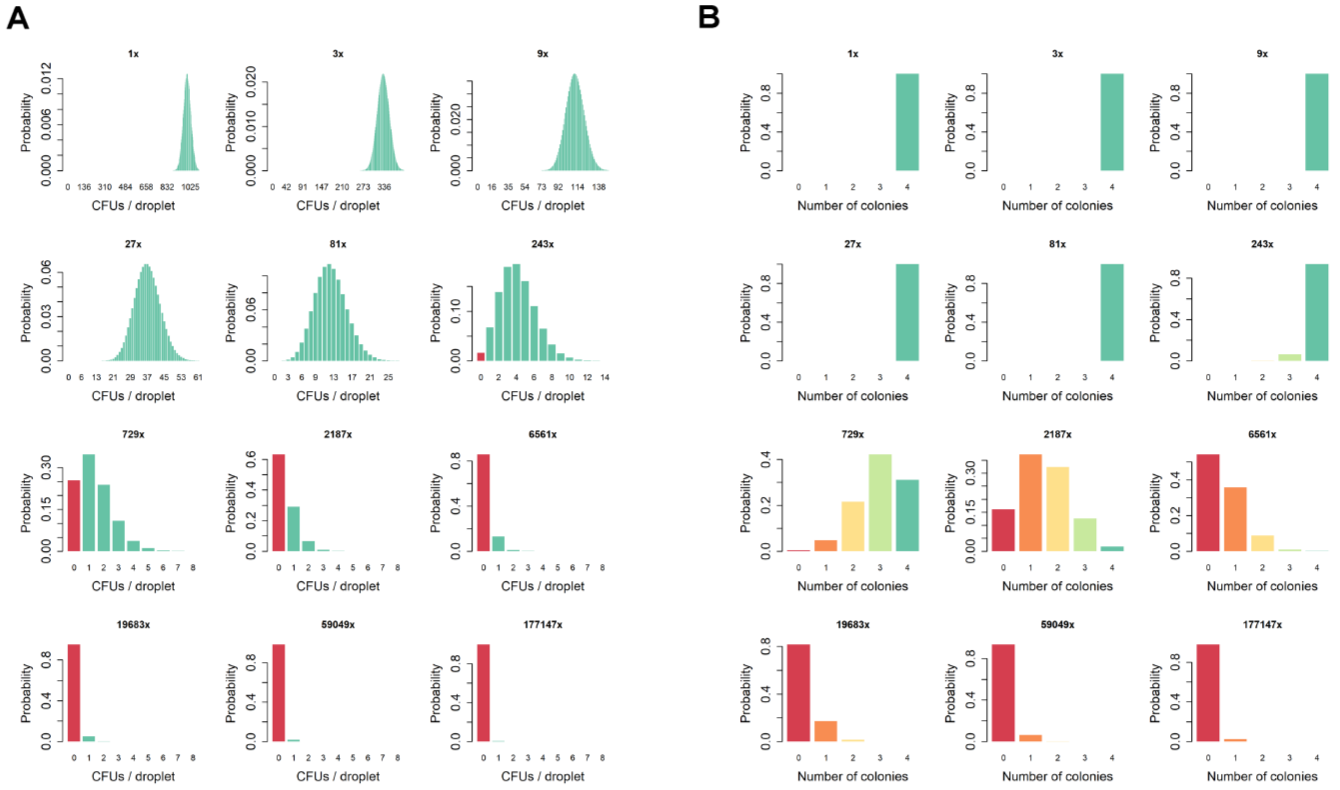


**Figure S3: Modelling of colony patterns produced by robotics-based CFU assay.**
Probabilities of observing different colony patterns are shown for the hypothetical case that a culture has 1000 CFUs / droplet, is serially diluted 3x across the plate, and then pinned in quadruplicate. Dilution factors are indicated above each bar plot.

**A.** The number of CFUs pinned is modelled as Poisson distributed. Bars are coloured green in the case that at least 1 CFU is pinned (i.e. there is a colony) and red in the case that no CFUs are pinned. At low dilution factors, there are still on average many CFUs / droplet, so it is highly probable that at least 1 CFU will be pinned and a colony will grow. As the dilution factor increases, the probability of no colony growing increases as the average number of CFUs / droplet decreases.

**B.** The number of colonies which grow at each dilution factor is modelling as binomially distributed, according to the probability of observing a colony from the Poisson distribution. Bars are coloured from green through amber to red according to the number of colonies which grow. At low dilution factors, it is highly probable that each position on the agar plate will contain a colony. As the dilution factor increases, the probability of not observing a colony increases, and hence it becomes more probable that less colonies will grow at these dilution factors.

A

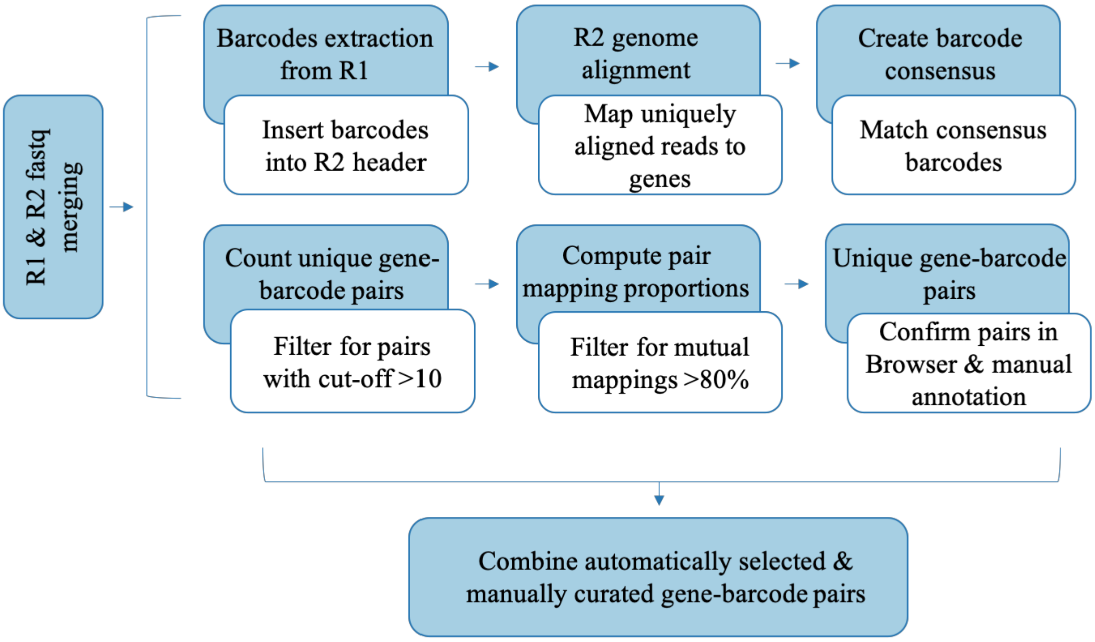


B

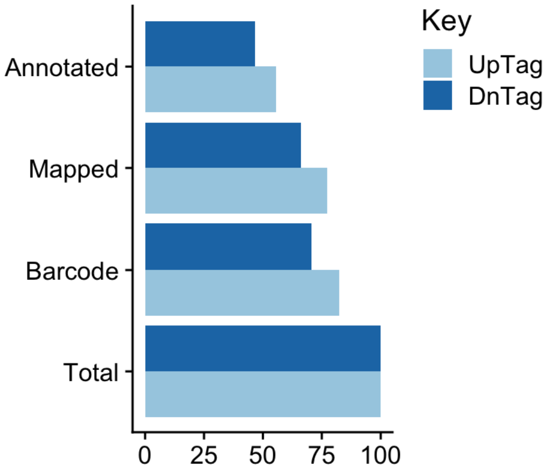


**Figure S4: Characterisation of deletion library.**
**A.** Scheme for deletion-library characterization using our in-house package, *BarSeqTools* (https://github.com/Catalina37/BarcountBarseqToolsPipelines/tree/master/BarseqTools). The main steps are as follows. The barcodes were extracted from R1 (forward reads) and assembled into consensus barcodes. Genomic sequences were extracted from R2 (reverse reads), aligned to the genome, and neighbouring genes identified. A table listing barcode-gene pairs assembled from R1 and R2 was generated. A gene was considered to be decoded (matched to a barcode) if: (a) the barcode-gene pair was observed frequently, i.e. at least 10 instances of that gene-barcode pair, (b) the gene was specific to that barcode, i.e. at least 80% of the reads from that gene map to that barcode, (c) the barcode was specific to that gene, i.e. at least 80% of the reads from that barcode map to that gene. Gene-barcode pairs were manually inspected using an in-house gene browser.
**B.** Reads loss following major analysis steps is recorded as a percentage of total reads (**Total**). Steps at which read loss was encountered includes read processing for barcode extraction **(Barcode)**, genomic sequence alignment to the reference genome with mapping of uniquely aligned reads **(Mapped)**, and mapping of these reads to the deletion library genes **(Annotated)**. Data are shown for both UpTag (bright blue) and DnTag (dark blue) barcodes.

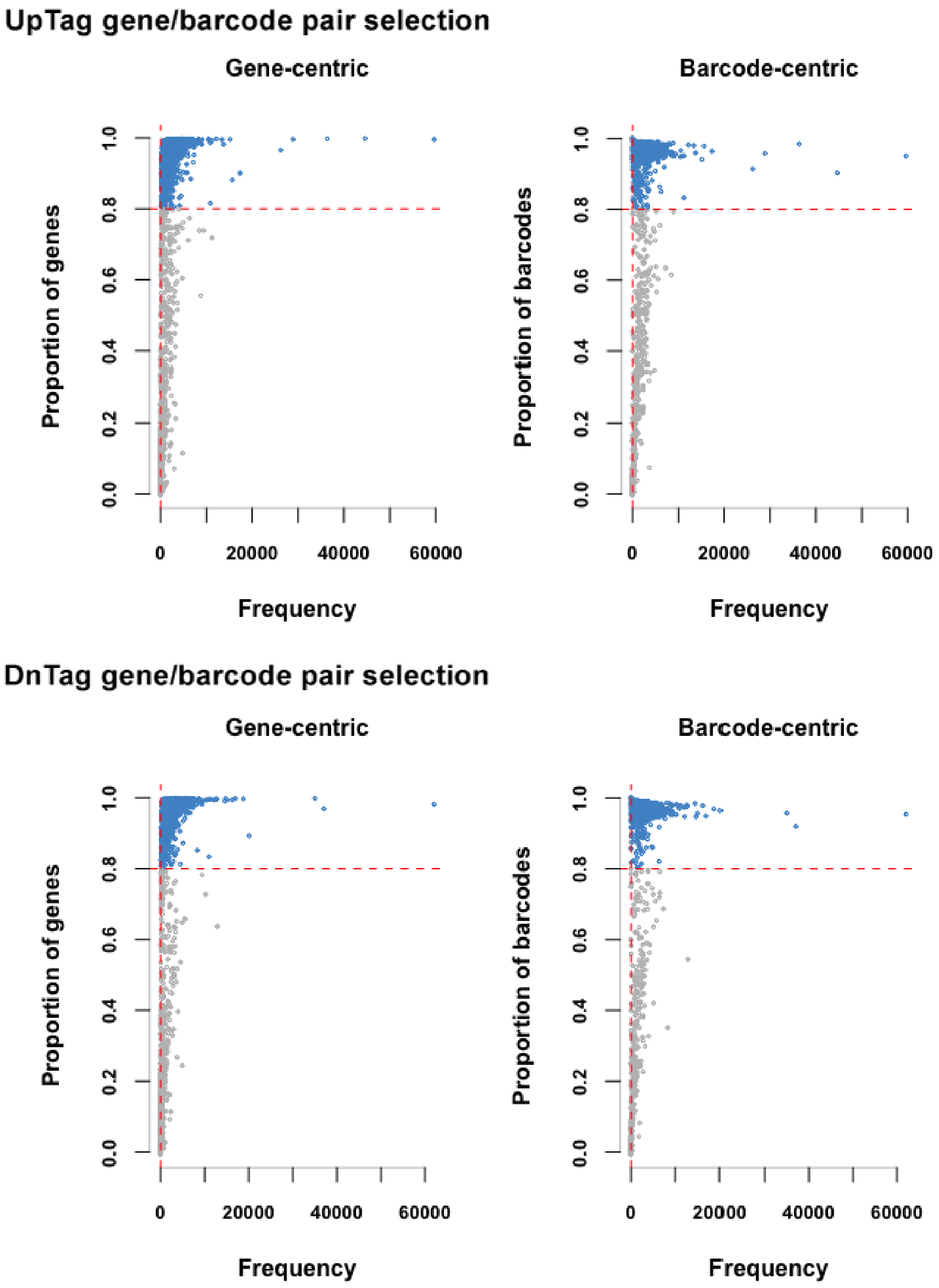


**Figure S5: Selection of high-confidence unique gene-barcode pairs.**
Mutual proportion of genes (gene-centric) and barcodes (barcode-centric) for each unique gene-barcode pair was calculated using the customized *aggregateProportions* function of *BarSeqTools*. Only gene-barcode pairs with proportions >80% and frequency >10 were selected automatically. These can also be visualised in the *GeneBrowser*.
